# Experimental and mathematical models reveal that AMPK modulates circadian clock gene expression and period through NAD+-dependent regulation of *Bmal1*

**DOI:** 10.64898/2026.09.29.755034

**Authors:** Alan Vandenberghe, Pauline Delpierre, Raoul Torero-Ibad, Pierre Leclerc, Anthony Treizebre, Alexandre Berthier, Laurent Heliot, Hélène Duez, Bart Staels, Philippe Lefebvre, Alessandro Furlan, Marc Lefranc

## Abstract

The circadian clock allows living systems to anticipate environmental daily variations by scheduling cellular mechanisms according to the time of the day. The reciprocal interplay between metabolism and circadian clock is essential for energy homeostasis, and the disruption of metabolic clock inputs contributes to dysfunction of clocks in peripheral tissues. However, the molecular mechanisms through which the feeding/fasting cycle entrains peripheral clocks remain incompletely understood.

Here, we investigate experimentally and theoretically the role of AMP-dependent kinase (AMPK), a key fasting sensor, in metabolic regulation of the circadian clock. Using a luciferase reporter driven by the *Bmal1* promoter to monitor molecular clock activity in human U2OS cells, and thanks to a signal processing method based on the Hilbert transform to retrieve the instantaneous phase and period of circadian signals, we show that pharmacological activation of AMPK by AICAR significantly elevates *Bmal1* promoter activity and markedly lengthens the clock period in a dose-dependent manner. Conversely, the clock period is shortened by the pharmacological inhibition by SR18292 of Peroxisome proliferator-activated receptor gamma coactivator 1-alpha (PGC-1-alpha). Importantly, the action of AICAR is abolished by FK866, a nicotinamide phosphoribosyltransferase (NAMPT) inhibitor, demonstrating that AMPK-dependent modulation of *Bmal1* promoter activity requires NAD+ availability.

These experimental results are recapitulated by numerical simulations of our previously published mathematical model, where PGC-1-alpha plays a key role in mediating AMPK activity to the clock. Moreover, the comparison of experimentally measured and theoretically predicted phase response curves, following AMPK activation at different circadian phases, provides an additional validation of the model.

By integrating biological experiments with mathematical modeling, our results identify a key mechanism through which metabolic factors can entrain and regulate the circadian clock via NAD+-dependent AMPK-mediated regulation of *Bmal1* promoter activity.

## INTRODUCTION

Most living systems experience the alternation of days and nights and the resulting periodic changes in their environment. To anticipate these changes and orchestrate their physiology accordingly, they rely on a circadian clock that provides them with an internal measure of time, a network of genes and proteins regulating each other so as to generate biochemical oscillations with a period of about 24 hours (Dibner, Schibler, & Albrecht 2010).

To maintain its synchronization with astronomical time, the biological clock must be informed of the progression of the day/night cycle, typically through the modulation of a core clock parameter by environmental cues. This external forcing allows the clock to maintain a stable phase relationship with the external cycle. Thus, clock function relies not only on its core oscillator but also on the pathways that synchronize it, and whose dysregulation can lead to clock dysfunction. This may explain why the central clocks of many organisms seem to have evolved strategies to protect themselves from fluctuations in daylight caused by changing weather (Pfeuty et al. 2011; Pfeuty et al. 2012).

While the light/dark cycle is a major synchronizer (Dibner et al. 2010; Albrecht 2012), other diurnal rhythms are also essential to schedule physiological processes, especially in key metabolic organs such as the liver or the pancreas, which are faced with varying levels of energy inputs (food intake) and expenditure (body exercise) across the day/night cycle (Sahar et Sassone-Corsi 2012). Accordingly, the hepatic clock is primarily entrained by feeding/fasting (Fe/Fa) cycles: inverting the Fe/Fa cycle with respect to the light/dark cycle results in a phase shift of the liver clock by almost 12 hours (Damiola et al. 2000; Mukherji et al. 2015; Manella et al. 2021). More generally, perturbations in the Fe/Fa timing, or in intake quality, have been shown to alter clock behavior. For example, mice subjected to a high-fat diet (HFD) display clock oscillations that are dampened and phase-shifted (Kohsaka et al. 2007; Hatori et al. 2012; Eckel-Mahan et al. 2013), thus altering the perception of time. Similar observations have been made in mice fed during the normal rest, fasting phase (Mukherji et al. 2015). Importantly, metabolic disorders follow gradually these perturbations in the clock oscillations, suggesting a causal relationship (Eckel-Mahan et al. 2013).

It is thus essential to understand mechanisms through which the liver clock is entrained by Fe/Fa cycles, and how modifications of liver clock inputs affect normal clock operation. This may allow one to identify the cause of some pathological situations and to design chronotherapeutical protocols to restore normal clock operation in such cases (see, e.g., (Woller et al. 2016; Ballesta et al. 2017)).

However, the exact nature of the mechanisms synchronizing the liver clock and other peripheral clocks to Fe/Fa cycles is still incompletely understood.

Several hypotheses have been proposed in the literature. For example, Lamia et al evidenced an AMPK-dependent CRY degradation mechanism and observed that AMPK activation shifted the clock (Lamia et al. 2009). Besides, it has been proposed that insulin, through upregulation of PER protein translation (Crosby et al. 2019), or PI3K signalling (Fougeray et al. 2022), are major clock synchronizers. Moreover, the SIRT1 protein, another energetic sensor, has been shown to control PER2 stability (Asher et al. 2008) and to modulate CLOCK-BMAL1 activity (Nakahata et al. 2008; Bellet et al. 2013). On top of that, other systemic factors such as glucagon or free fatty acids (FFA) are known to influence clock genes and contribute to clock resetting (see, e.g., Mukherji et al. 2015). Anti-cancer translation inhibitors were shown to alter the molecular circadian clock through the downregulation of PER2 and NR1D1 levels (Berthier et al. 2023). The multiplicity of potential clock input pathways thus makes it difficult to evaluate their relative importance in the metabolic synchronization of the clock, especially as the core network itself consists of many interlocked feedback loops.

To disentangle this complexity and gain insight into the synchronization of the hepatic clock to metabolic rhythms, we previously designed a mathematical model incorporating additional feedback loops featuring SIRT1 and AMPK, two important physiological actors which respectively sense the key metabolites NAD+ and AMP (Woller et al. 2016). This model reproduced faithfully gene expression and NAD+ profiles from mouse livers, as well as the modified gene expression patterns observed in mice invalidated for the AMPK upstream regulatory kinase LKB1 (Lamia et al. 2009).

Importantly, this study pointed both to a possible role of the *Bmal1* gene as receiving metabolic input to the core clock and to the importance of the PGC-1-alpha protein as a clock actor, integrating NAD+ and AMP signaling (Cantó et al., 2010), and inducing *Bmal1* through coactivation of RORa and RORg (Liu et al., 2007). These predictions were experimentally and theoretically confirmed by Foteinou et al (2018) who showed that incorporating SIRT1 and PGC-1-alpha in their mathematical model was required to numerically reproduce their experimental observations in U2OS and NIH3T3 cells, and that silencing one or the other gene affected the clock (Foteinou et al. 2018). However, Foteinou et al. did not consider AMPK in their analysis, focusing on SIRT1 only. In contrast, Woller et al. proposed that AMPK was the main input, assuming that the daily variation of SIRT1 activity was mostly due to the NAMPT-driven NAD+ salvage pathway (Ramsay et al. 2009).

To further investigate this issue and obtain additional experimental evidence supporting or challenging the importance of the AMPK - PGC-1-alpha - BMAL1 axis in mammalian clock signaling, we sought to experimentally characterize the action of AMPK on the mammalian circadian clock. To this aim, we administered AICAR, an AMP analogue, to U2OS-B6 cells, a well-established human cell line for studying circadian oscillations, studying their response using a *Bmal1:luc* luminescent reporter stably integrated in the cell genome (Vollmers et al. 2008). We observed that AMPK activation significantly elevated the level of the reporter gene activity and that it induced a systematic increase of the period with increasing AICAR dose. Moreover, the clock period was significantly reduced upon administration of SR18292, a pharmacological inhibitor of PGC-1-alpha. Remarkably, the action of AICAR was abolished when the NAMPT inhibitor FK866 was simultaneously delivered with AICAR, blocking NAD+ recycling. Finally, we constructed an experimental phase response curve (PRC), which indicates for the first time the phase shift experienced by the U2OS clock upon AMPK activation by AICAR at different times of the cycle.

Using a marginally adapted Woller model, where the dominant input is the AMPK-PGC-1-alpha - BMAL1 axis, we were able to reproduce quantitatively the elevated *Bmal1* transcription and the variation of the clock period with AICAR dose. The general structure of the PRC measured was also well reproduced with our model.

Our study thus represents a significant step towards a systematic characterization of the mechanisms through which AMPK acts on the circadian clock. Our findings are globally consistent with the predictions by Woller et al. (2016) and their hypothesis that the AMPK-PGC-1-alpha - BMAL1 axis is a strong metabolic input to the clock. They identify AMPK as a major input of the clock, not only strongly modulating *Bmal1* expression levels but also resetting significantly the clock phase.

## RESULTS

### Cell circadian clock characterization based on the monitoring of *Bmal1* promoter activity

Luminescent-based reporters are precious tools to study molecular mechanisms over time. We thus used this approach to monitor the circadian clock, by taking advantage of U20S cells stably expressing destabilized luciferase under the control of the *Bmal1* promoter (-422 to +108) (Vollmers et al. 2008), thereafter called U20S-B6 (Fig.1A).

**Figure 1.**
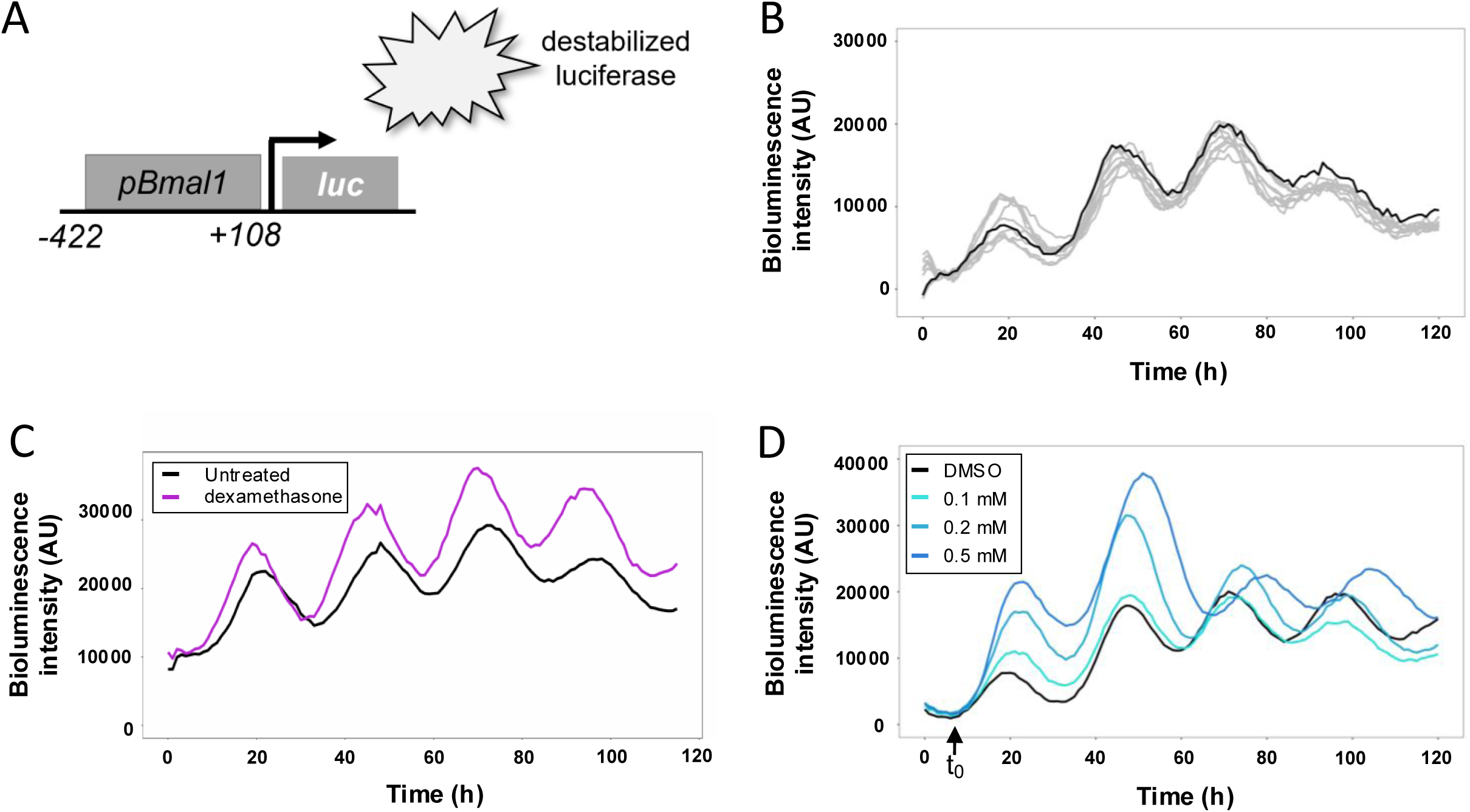
Quantifying *pBmal1* activity to monitor circadian clock over time. A. A circadian reporter based on the expression of destabilized luciferase under the control of bmal1 promoter is stably expressed in U2OS B6 cells. B. Raw signal measured in U2OS B6 cells. Individual signals from untreated condition are represented in grey and the mean signal in black. n=9. C. Synchronization of U2OS B6 cells with dexamethasone applied during 2h at 100 nM (purple) was compared to the simple medium refreshment in control condition (black). D. Raw signals measured in U2OS B6 cells. Signals from DMSO control are represented in black, while blue curves correspond to cells treated with increasing concentration (0.1, 0.2 and 0.5 mM) of AICAR. The t

Using this model, we assessed *Bmal1* promoter activity in cells over an extended period of time (5 days) by measuring the luminescence level every 30 minutes with a plate reader equipped with temperature and CO_2_ modules. The number of cells was adjusted to 20,000 cells per well in a 96-well plate in order to obtain a signal high enough, yet not saturated, with respect to the sensitivity of the detector. This allowed us to directly observe circadian oscillations in *pBmal1* promoter activity (Fig.1B).

Noteworthy, curves from single wells in a single experiment were very similar one to another, demonstrating the robustness of the approach, and peaks from different experiments were very synchronous, suggesting that medium replacement to integrate luciferin at the beginning of the experiment is sufficient to synchronize U2OS-B6 cells. Although dexamethasone, a classical clock synchronizer, could increase the average level and amplitude of oscillations (Fig.1C), its use did not drastically modify the luciferase cycling pattern. We therefore chose not to use this glucocorticoid further in our experiments, in order to avoid any additional interference with the molecular clock.

Our previously published mathematical model highlighted a key role for AMPK and SIRT1 inputs, integrated by the PGC-1-alpha protein to regulate the clock as a function of energetic status (Woller et al. 2016). We decided to investigate further this concept by modulating AMPK activity in U20S-B6 cells using the pharmacological activator AICAR (5-amino-4-imidazolecarboxamide ribonucleoside), an AMP analogue. We first determined the appropriate AICAR concentration to elicit responses in these cells without affecting cell viability. For that purpose, we incubated cells with AICAR concentrations ranging from 0.1 to 2 mM and monitored them for 5 days (Fig.S1).

Since medium change had been observed to reset the clock significantly, as previously reported in the literature (Welsh et al. 2004; Guenthner et al. 2014), it was critical not to wash out AICAR to avoid disturbances on clock phase measurement. Because of the possible toxic effects of AICAR, this forced us to assess the AICAR levels that cells could withstand across an extended period of time. When compared to untreated U20S cultures, AICAR concentrations up to 0.5mM did not have any effect on cell confluence over time. Cells treated with 1 mM AICAR grew more slowly, and cells treated with 1.5 and 2 mM did not grow and died over the 5-day treatment period (Fig. S1).

We then used a validated genetically encoded FRET biosensor, namely AMPKAR-EV (Konagaya et al. 2017), with which AMPK activation was monitored over time in response to 0.1 to 1 mM AICAR concentrations. We observed a dose-dependent AMPK activation, with 0.5m M AICAR leading to a near-optimal activation rate, as previously described in the literature (Gowans et al. 2013) and close to that observed with short-term 1 mM AICAR or 2-deoxyglucose, which were used as positive controls (Fig. S2). Thus, we chose 0.5 mM as the maximal concentration for our five-day experiments.

### AMPK activation induces *Bmal1* promoter activity as indicated by increased reporter luminescence

Subsequently, we administered 0.1 to 0.5 mM AICAR to U20S-B6 cells and monitored the promoter activity over a period of five days. Strikingly, AICAR significantly impacted circadian oscillations in a dose-dependent manner, with a mean luminescence intensity significantly higher in treated compared to control cells, and a shift in time clearly observable with 0.5 mM AICAR (Fig.1D).

The significant increase in reporter signal pointed to a strong dependence of *Bmal1* transcription on AMPK activity, establishing the latter as a major clock input. We found that the maximal effect of AICAR on promoter activity level was reached during the 2^nd^ circadian cycle, displaying approximately a 2-fold increase with 0.5 mM AICAR.

A notable observation is that the signals for the different AICAR doses are superimposed throughout the early stage of the experiments (Fig. 1 D), despite the different AICAR treatments. This uniformity in the response cannot be attributed to a delay between AICAR administration and its action on the clock, as Fig. S2 shows that cells respond to AICAR within minutes. Thus, there exists a window of insensitivity of *Bmal1* promoter activity to AMPK activation in a time window around the first minimum of the luminescence signals, which is observed about 7 hours after the medium change in which AICAR is delivered. Here and below, we used this first minimum (shown as t_0_ in Fig.1D) to define the origin of the circadian cycle. Since the half-life of the luciferase used is short (0.4 h), we assumed that the luminescence signal closely follows *Bmal1* transcriptional activity and that beginning of the cycle thus defined approximately coincides with minimal *Bmal1* promoter activity.

To make this analysis more quantitative, we went on to analyze the time derivative of the luminescence signals, which reflects the balance between synthesis and inactivation/degradation of luciferase and allows us to monitor changes in these two processes. Because numerical differentiation is sensitive to noise, we first applied a basic exponential moving average filter to the raw signals of Fig. 1D, after aligning their levels at time t0 to account for a small variability in cell count between wells (Fig. 2A). The time constant of the filter was about 2.5 hours, but we checked that results did not depend on it, except for the smoothness of the curves. We then estimated the time derivative of the signal with a simple order-2 finite-difference scheme (Fig. 2B). Finally, we plotted the difference between the derivative estimate for each AICAR dose and the one obtained for the control curve (Fig. 2C), the deviations from 0 thus indicating the effect of AICAR administration.

**Figure 2.**
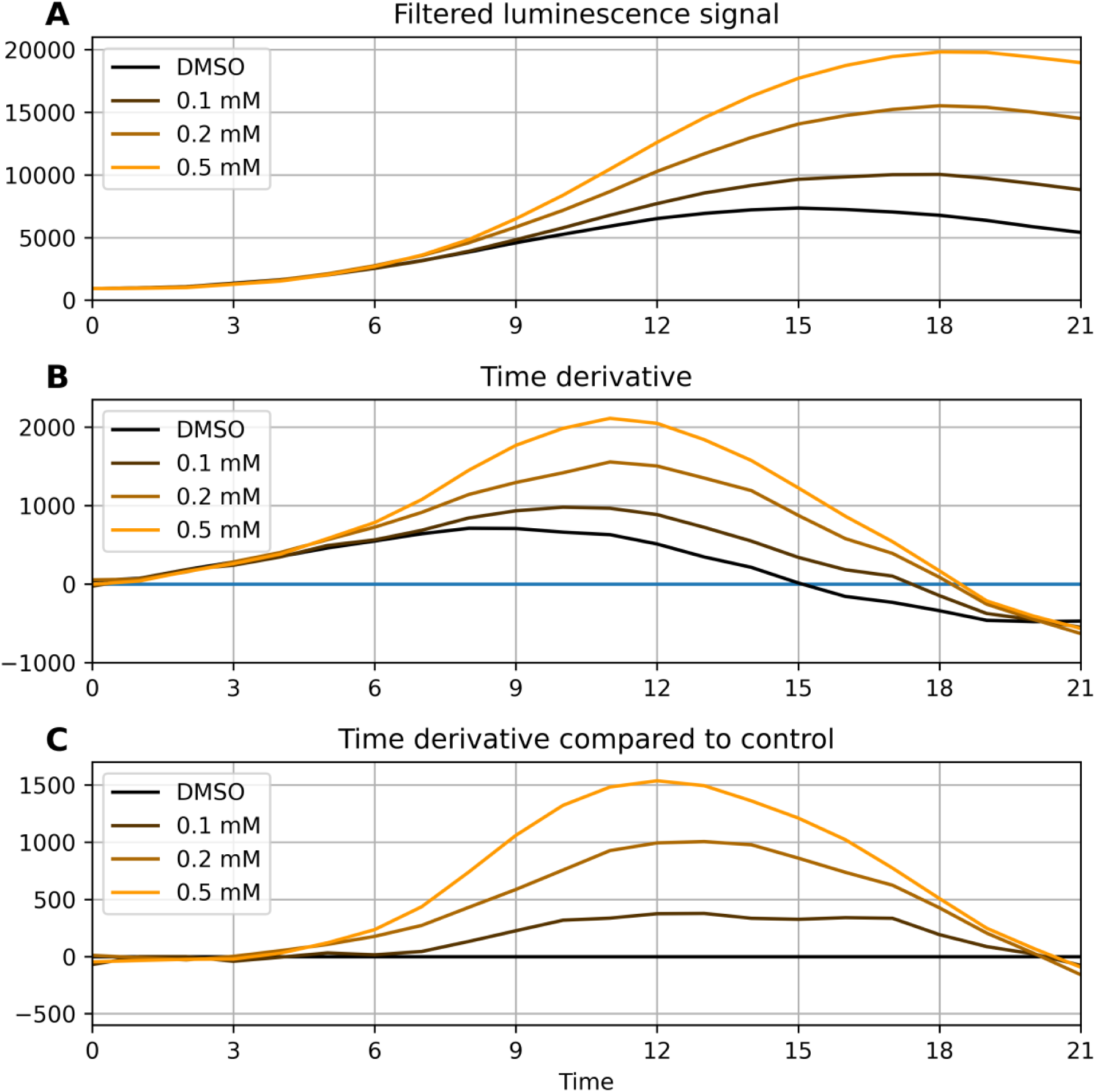
Dynamical analysis of luminescence signals. A. The luminescence signals from Fig.1D have been processed through an exponential moving average filter with time constant 2.5 h to prepare them for the subsequent numerical differentiation. Time 0 represents the origin of the cycle (marked as t_0 in Fig. 1D) and corresponds to minimal Bmal1 promoter activity. B. Time derivative of the luminescence signals obtained with a simple order-2 finite-difference differentiation scheme. C. Difference between the derivative estimates for the different doses with the estimate of the control curve (vehicle only).

The curves in Fig. 2C show that at all doses tested, AICAR has no detectable effect during the first 4-5 hours of the cycle, consistent with the superimposition of the luminescence signals and their derivatives (Figs. 2A-B). The effect then gradually increases, reaching a maximum approximately 12 hours after the beginning of the cycle. Beyond this point, however, the analysis becomes difficult as signals obtained at different doses display different interpeak intervals. Thus, values at the same time do not correspond to the same phase of the circadian cycle.

This observation strongly suggests that AMPK does not stimulate *Bmal1* transcription directly but through a positive transcription factor required for *Bmal1* transcription, the effect vanishing in the absence of this transcription factor and being maximal when it is strongly bound.

### A novel workflow to describe clock features, and to better characterize AMPK action on it

With the intent to characterize optimally these circadian oscillations and to obtain subsequently a quantitative assessment of their modulation by metabolic inputs, we set up a novel workflow aiming at measuring accurately the instantaneous phase of the luminescence signals. For that purpose, we first processed the raw signals using a band-pass order-2 Butterworth filter from the SciPy library (Virtanen et al. 2020). Eliminating low-frequency components allowed us to get rid of long-term variations in the setup, including variations in cell count, without assuming a specific law of variation, while the short-term fluctuations were cleaned by removing high-frequency components. The frequency window used for the Butterworth filter was fixed by examining the frequency spectrum obtained with the Fast Fourier Transform, and ensured that periodic components corresponding to periods between 16 and 36 hours are preserved (Fig. 3A).

**Figure 3.**
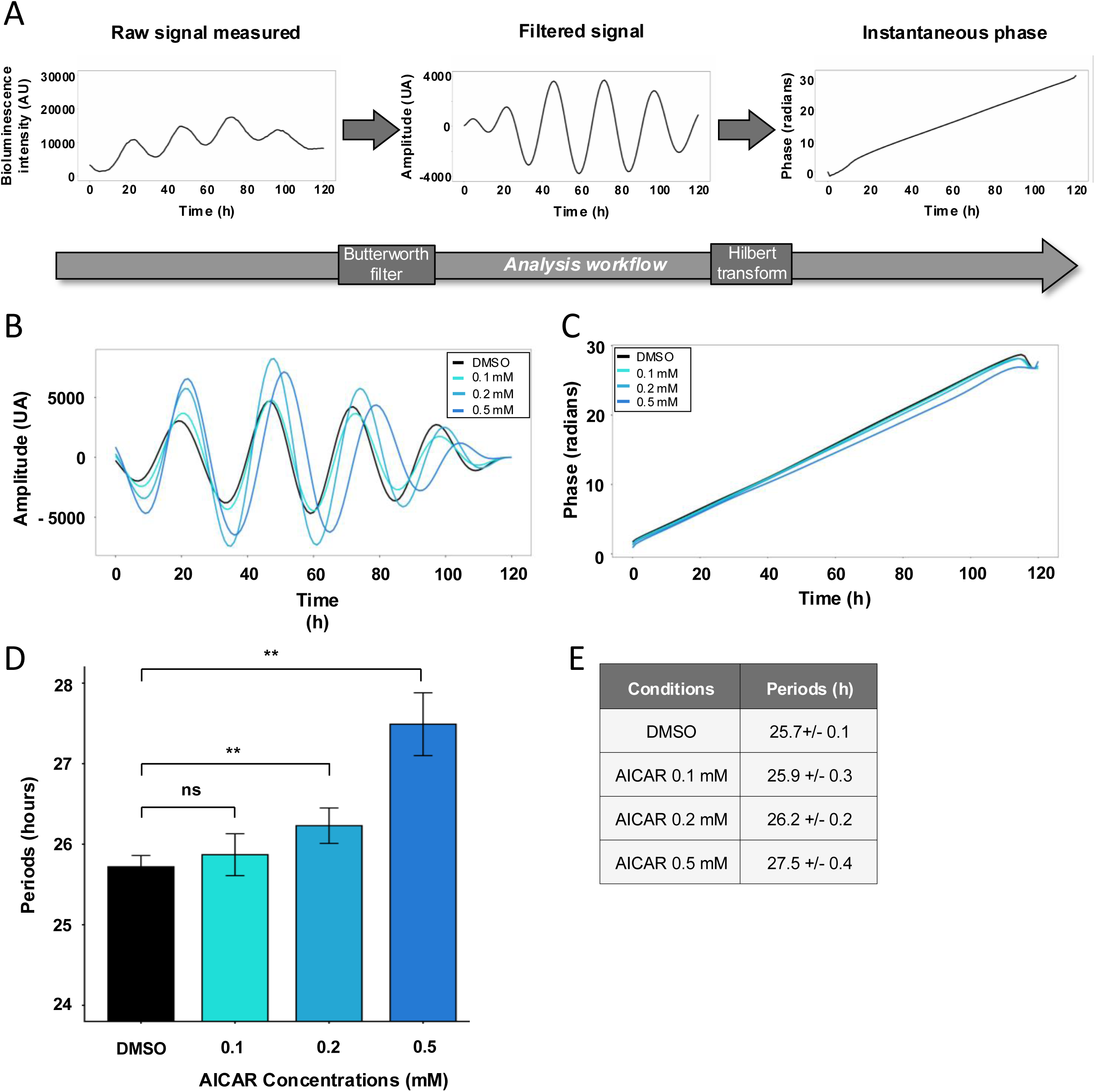
A novel workflow to better characterize cell circadian clock, and the impact of AICAR on it. A. An analysis workflow was set up to treat the oscillating signal with Butterworth filter and Hilbert transform in order to calculate phase and period of this signal. B Filtered signals from raw data of cells treated with AICAR. Blue curves represent increasing concentration (0.1, 0.2 and 0.5 mM) of AICAR while the black curve represents the DMSO control condition. C. Hilbert transformed was applied to filtered signals in B to calculated instantaneous phases. D. Periods were inferred from the instantaneous phases in C during the interval 40-110h of the experiment, There are represented in blue shades for increasing concentrations (0.1, 0.2 and 0.5 mM) of AICAR, and in black for DMSO cells. Statistical test was performed between AICAR conditions and DMSO condition. Statistical tests were performed with Wilcoxon-Mann-Whitney test for comparisons, and p-values adjusted using the Holm correction, ns: non significant, **: p<0.01. N=3 independent experiments, n=9. E. Table of periods in hours +/-SD.

Thus, we obtained a signal with a clear circadian oscillatory component, which makes the determination of the instantaneous signal phase easier. Obviously, the period of the signal and the phase difference between two signals are left unchanged by the linear filtering, which is adequate for our characterization of the circadian clock. Using the Hilbert transform, the oscillatory signal was then transformed into the so-called analytical signal, which rotates in the complex plane, and from which amplitude and phase can easily be computed as being the modulus and argument of this complex signal.

The instantaneous phase displayed a very linear asymptotic behavior (Fig.3A), with its slope providing an accurate estimate of the clock period using a linear regression, more precisely than by computing interpeak intervals. The remarkable constancy of the slope (except near the very end, due to numerical boundary effects) suggests that there was no significant depletion in nutrient availability or AICAR activity through time. Subsequently, the period of circadian oscillations was quantified to last 25.7±0.2 h in vehicle-treated U20S-B6 cells.

To assess precisely the dependence of the clock period on AMPK dose, luminescence raw signals were then analyzed according to the afore-described workflow, with Butterworth filtering nicely evidencing a growing delay in the oscillations when the AICAR dose delivered to cells increased from 0.1 to 0.5 mM (Fig.3B). The oscillation periods were then calculated from the variation of the instantaneous phases (Fig.3C) with time, using a linear regression. Administrations of AICAR at 0.1, 0.2 and 0.5 mM concentrations increased this period from 25.7± 0.1 h for vehicle-treated cells to 25.9±0.3 h, 26.2±0.2 h, and 27.5±0.4 h respectively (Fig.3D-E). It also must be noted that, in comparison, pretreatment with the gold standard dexamethasone reduced the period only marginally, from 25.7 to 25.6 h.

### A minimally modified Woller model taking into account the nuclear translocation of activated PGC-1-alpha nicely fits the experimental observations

We then sought to determine whether the important increase in *Bmal1* transcription and the period lengthening triggered by AICAR administration were consistent with predictions of the mathematical model that we previously established for hepatocytes⍰⍰. Although hepatocytes differ substantially from U2OS cells physiologically, a key ingredient of the Woller model, namely the AMPK-PGC-1-alpha-BMAL1 axis, may nevertheless operate similarly in the two cell types. It was thus interesting to test whether the observed increase in the period could be reproduced by this model.

A first step was to determine if we could reproduce the significant increase in the average *Bmal1* expression observed upon AMPK activation with 0.5 mM AICAR. Interestingly, we found that merely increasing AMPK activity in the model was not sufficient, as it could only induce a 25% increase of *Bmal1* expression. We thus carried out a sensitivity analysis of the Woller model, varying all kinetic constants in turn to identify those that control *Bmal1* expression and could therefore be considered as putative targets of AMPK activation. The result of this systematic analysis is shown in

Fig. 4A, where for each kinetic constant, we have plotted the *Bmal1* expression temporal profile with the largest amplitude observed as the value of the constant was gradually varied between 20% and 500 % of its nominal value. Remarkably, we found that a 2-fold increase in peak *Bmal1* transcription could be achieved by varying a single kinetic constant only through two mechanisms, acting directly on *Bmal1* and not on other actors: modulation of the basal Bmal1 transcription rate, either directly or indirectly via the abundance of nuclear PGC-1-alpha, or modulation of Bmal1 mRNA stability (Fig. 4A). In particular, the peak *Bmal1* mRNA level is almost insensitive to the kinetics of its repressor NR1D1.

**Figure 4.**
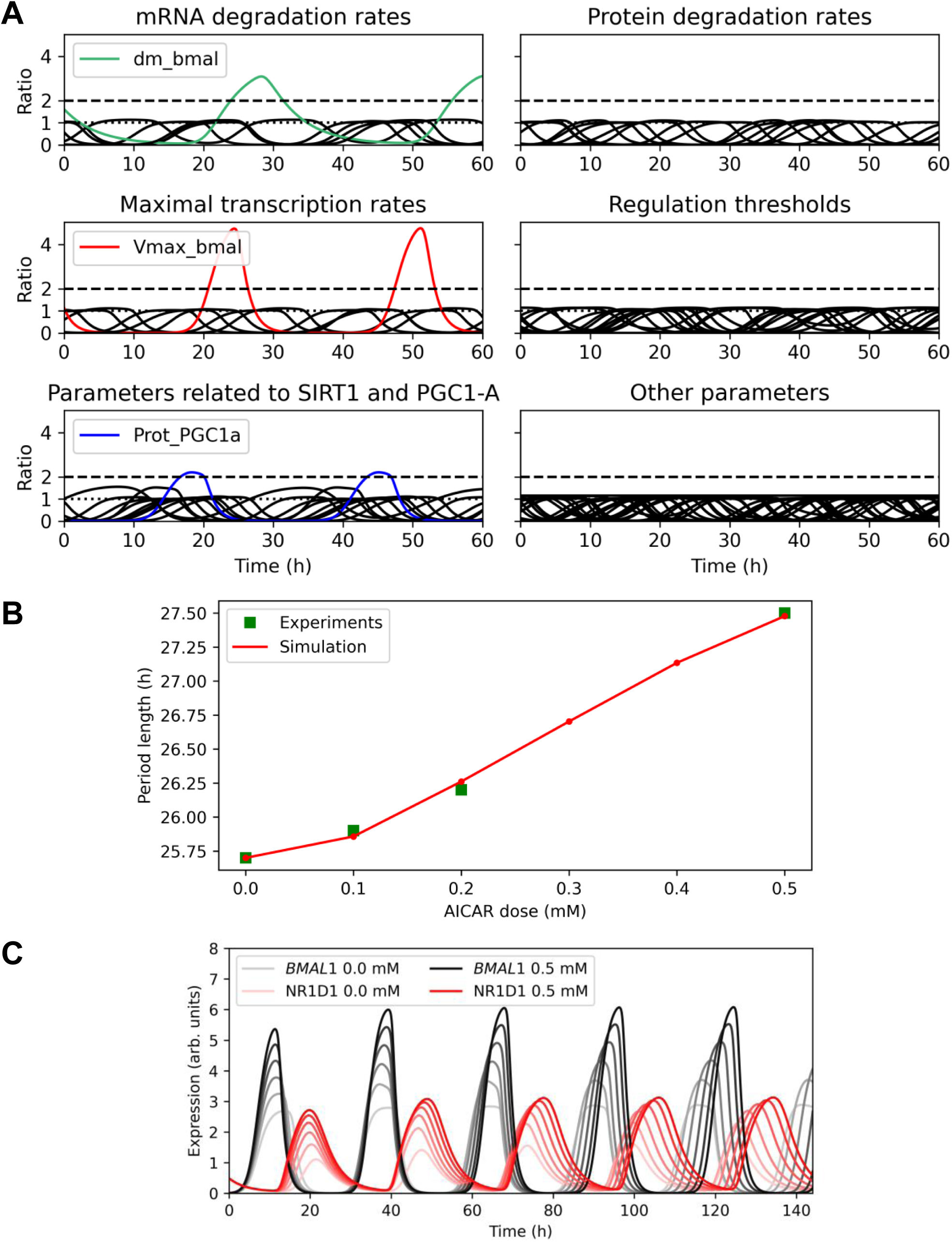
Getting insight into the observed clock behavior using the Woller model. A. Each kinetic constant of the Woller model was changed gradually from 20% to 500% of its nominal value, storing the temporal profile with the highest increase in *Bmal1* amplitude. The corresponding *Bmal1* time profiles are displayed, grouped according to the type of kinetic constant varied. The 2-fold increase in *Bmal1* transcription observed experimentally (indicated by a dashed line) can only be reproduced by enhancing directly the basal *Bmal1* transcription rate or PGC1-A abundance and activity. B. After the Woller model is rescaled to match the zero-dose U2OS-B6 period, the kinetic constants describing the variation of AMPK activity and PGC1-A nuclear abundance with AICAR dose are adjusted to reproduce the observed dependence of the period with AICAR dose. All other kinetic constants of the Woller model are left unchanged. C. The temporal profiles for the *Bmal1* expression and NR1D1 abundance profiles are shown for different AICAR doses. The increased NR1D1 amplitude that results from highest *Bmal1* expression contributes to period lengthening by increasing the delay between two *Bmal1* peaks.

To refine the mathematical model so that it reproduces the elevated *Bmal1* promoter activity upon AMPK activation, we examined whether one of the three identified constants could be made AMPK-dependent. We found that stabilizing *Bmal1* mRNA while activating AMPK suppressed oscillations. Since PGC-1-alpha nuclear abundance directly affects *Bmal1* transcription, the most parsimonious hypothesis was to assume that it depends on AMPK activity. In fact, the above analysis revealed a shortcoming of the Woller model, which considers activation of PGC-1-alpha, but does not describe its AMPK-dependent nuclear translocation (Anderson et al. 2008). Remarkably, simulations of a Woller model modified so that the PGC-1-alpha nuclear abundance depends linearly on AICAR dose easily reproduced the significant experimental increase in *Bmal1* promoter activity observed upon AICAR delivery.

To compare the model predictions with the other experimental results, we treated basal AMPK activation and PGC-1-alpha nuclear abundance, together with the slopes of their dose-response relationships to AICAR, as adjustable parameters. This is reasonable given the many differences in physiological context between hepatocytes and U2OS-B6 cells and because the relation between AICAR dose and AMPK activation was not investigated by Woller et al. (2016). While many other kinetic parameters could plausibly differ between the two situations, it was interesting to test a minimally adapted mathematical model against experimental data to assess its robustness to differences between cellular phenotypes. We also rescaled the model time so that for vehicle-treated U2OS-B6 cells, the theoretical period matches the experimental period measured, allowing us to test whether the model predicted correctly the relative changes in period length as AICAR dose is increased.

By fitting the four above-mentioned kinetic constants, we were able to obtain a perfect agreement between the experimentally measured periods and those predicted by the mathematical model while enforcing a 2-fold increase in maximum *Bmal1* mRNA expression (Fig. 4B).

Then, by carrying out numerical simulations of the Woller model for different AICAR doses between 0.0 and 0.5 mM, we studied more closely the gradual period lengthening observed upon increasing the AICAR dose, besides a clear increase in *Bmal1* expression level (Fig.4C). As *Bmal1* expression is strongly increased due to enhanced PGC-1-alpha activity, *NR1D1* expression is also significantly boosted. Higher abundances for the NR1D1 protein imply longer decay times and thus longer time intervals during which *Bmal1* expression is repressed, leading to the period lengthening observed. These numerical simulations of the model therefore confirm that the period lengthening observed for increasing AICAR doses is globally consistent with the elevated expression of *Bmal1* and likely results from it.

### PGC1a inhibition by SR18292 shortens the clock period

According to the literature and to our mathematical model, AMPK can act on *Bmal1* transcription through the phosphorylation-dependent activation of PGC-1-alpha. This phosphorylation is a prerequisite for the subsequent deacetylation of PGC-1-alpha by SIRT1, required for the optimal activity of PGC-1-alpha as a co-factor of *Bmal1* transcription (Cantó, 2010). Therefore, we then investigated whether pharmacological inhibition of PGC-1-alpha would trigger a reverse impact on the clock period.

For that purpose, we administered to U2OS-B6 cells increasing doses of the PGC-1-alpha antagonist SR18292 (Sharabi et al. 2017), from 5 µM to 20 µM (higher doses becoming toxic along the 5-day incubation. SR18292 catalyzes the acetylation of PGC-1-alpha, thus deactivating it. Interestingly, SR18292 administered at 20 µM markedly shortened the clock period to 22.5 h (Figs. 5A-D). Interestingly, this also induced a strong attenuation of *Bmal1* reporter oscillations (Fig.5A). To better illustrate the effect, the amplitudes of the oscillations were obtained as the moduli of the analytical signal for different SR18292 doses and are shown in Fig. 5E. There is a clear decrease in the amplitude starting at 40-50 hours following treatment. We do not know whether this is due to a collapse of individual cellular clocks or to a desynchronization of the different cells; however, this points to an important perturbation of the clock.

**Figure 5.**
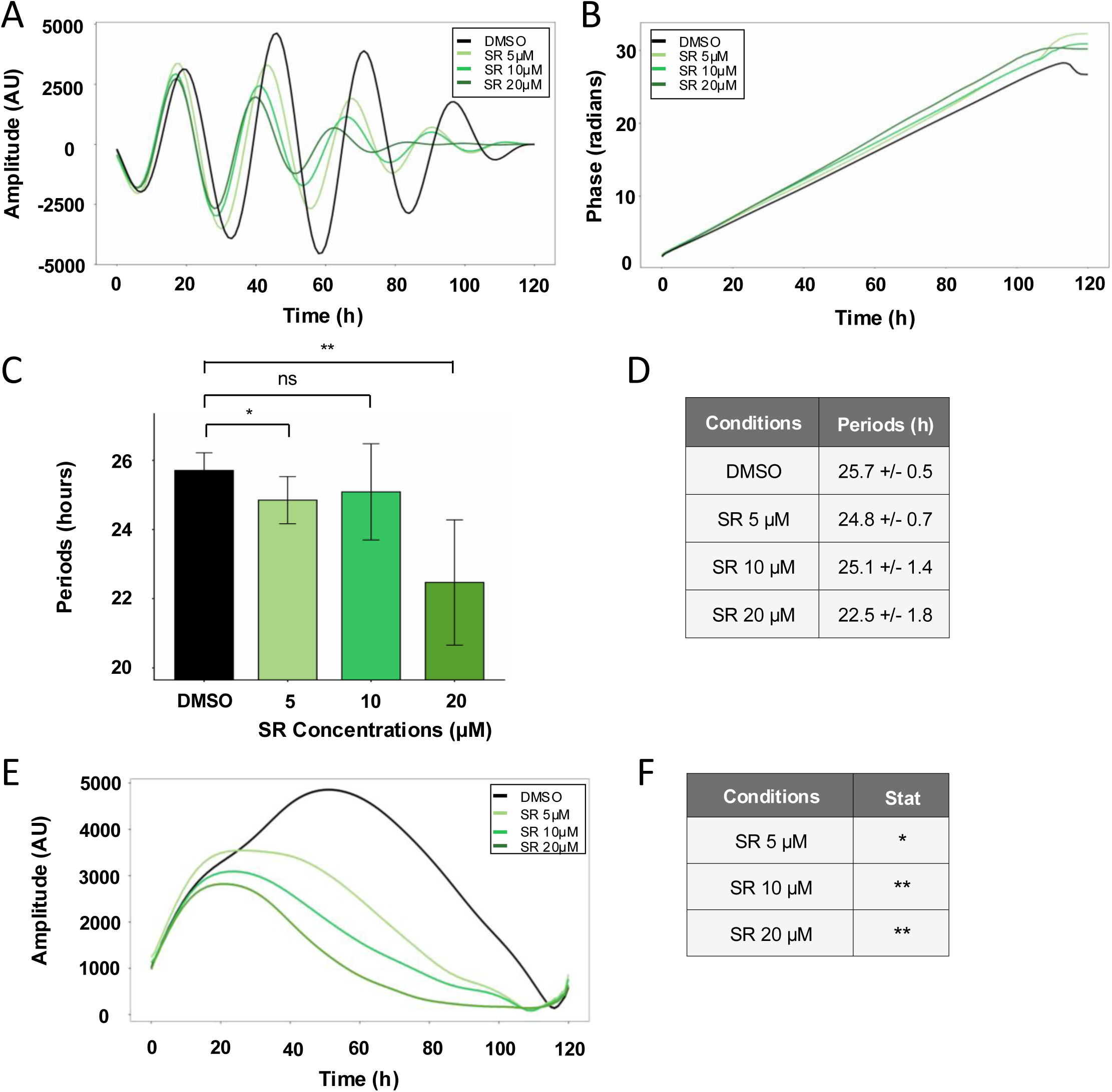
Effect of SR18292 on *Bmal1* promotor activity. A. Filtered signals processed using a Butterworth filter are shown in green for SR treatment at 5 µM, 10 µM, and 20 µM, with the DMSO control shown in black. B. Instantaneous phases calculated from A using Hilbert transform. C. Periods were inferred from the instantaneous phases in B during the interval 40-70h of the experiment. Statistical tests were performed with Wilcoxon-Mann-Whitney test for comparisons, and p-values adjusted using the Holm correction, ns: non significant, *: p<0.05, **: p<0.01. N=3 for 3 independent experiment, n=9. D. Table of periods in hours +/-SD. E. Instantaneous magnitude calculated from A using Hilbert transform. F. Comparison with DMSO using statistical test applied on area under curves (AUC) calculated from E, and performed with Wilcoxon-Mann-Whitney test for comparisons. p-values are adjusted using the Holm correction, ns: non significant, *: p<0.05, **: p<0.01. N=3 for 3 independent experiment, n=9.

We then tried to combine the AICAR and SR18292 treatments to assess whether a pharmacological inhibition of PGC-1-alpha would neutralize the action of AICAR. We used 0.5 mM AICAR and 20 µM SR18292, the doses that induced the clearest effects when used alone. The result was difficult to interpret due to a non-stationary behavior of the clock. This is illustrated by the dynamical behavior shown in Fig. S4A: the AICAR+SR18292 curve is well ahead of the DMSO control at the second peak, suggesting a significantly shorter period, but gradually lags the control thereafter, consistent with a longer period. Consequently, the overall phase variation across the experiment was comparable between the DMSO and AICAR+SR18292 conditions (Fig. S4A). To quantify this effect, we used the instantaneous phase curves of Fig. S4B to compute the phase advance over 26 hours and convert it into an equivalent period length (Fig. S4C). This showed that the effect of SR18292 combined with AICAR was initially comparable to that observed when the inhibitor was used alone, inducing a significant period shortening, but then became very similar to that of AICAR. Indeed, over the 40-100 h interval, the Hilbert transform analysis yielded similar periods for the AICAR and AICAR+SR18292 conditions (Fig. S4D). These observations are consistent with a progressive attenuation of the SR18292 effect by AICAR. Moreover, the simultaneous administration of SR18292 and AICAR strongly attenuated the oscillation amplitude (Figs. S4E–F), similarly to treatment with SR18292 alone (Figs. 5E–F).

We reasoned that these nonlinear effects could result from AMPK activation increasing NAD+ levels and thereby promoting PGC1-alpha deacetylation by SIRT1, as described by Cantó et al (2009), potentially interfering with SR18292-mediated acetylation. Interestingly, we could not identify any study in the literature in which both compounds had been used simultaneously in the same cells. Based on this hypothesis, we focused on NAD+ and investigated the impact of its depletion on the clock in our system to obtain further insights on AMPK action.

### NAMPT inhibition interferes with AMPK-driven clock period lengthening and *Bmal1* **promoter activation.**

SIRT1 is a NAD^+^-dependent deacetylase which is key to the activation of PGC-1-alpha, and maintenance of the NAD^+^ pool in cells strongly relies on a salvage pathway controlled by NAMPT (Ramsay et al. 2009). To interfere with this activity and subsequently investigate the impact of AICAR on the clock period length, cells were first incubated with the pharmacological NAMPT inhibitor FK866. As with AICAR, we initially aimed at defining the adequate subtoxic FK866 concentration to use. We scanned a range encompassing concentrations from 1 to 20 nM, according to doses classically used in the literature. This allowed us to conclude that FK866 was well tolerated by U2OS-B6 cells up to a concentration of 5 nM, before becoming toxic for cells at 10 nM (Fig. S5). With subtoxic concentrations, FK866 hardly affected the clock period length (Fig. S6).

To investigate whether FK866 action could inhibit the effect of AICAR on *Bmal1* transcription, based on the hypothesis that AMPK-driven phosphorylated PGC-1-alpha requires NAD+-dependent SIRT1 activity to act on p*Bmal1*, we combined the administration of both pharmacological agents. We started with 5 nM FK866 and 0.5 mM AICAR concentrations, but these combinations were unfortunately toxic to cells, as evidenced by cell observation and luminescence signals close to the noise levels. Interestingly, reducing FK866 concentration to 2.5 nM in such combinations allowed us to monitor cell populations over the five days of the experiment, and to evidence that FK866 greatly reduced AICAR-driven clock period increase with 0.5 mM AICAR concentrations (Fig.6A and 6B; raw data in Fig.S7). Indeed, the clock period length was then reduced from 27.8+/-1.7 hours with AICAR alone to 26.0 ± 0.2 hours with the combination, close to the period length for control U2OS-B6 cells, indicating that the action of AMPK on the circadian clock is strongly NAD+-dependent.

**Figure 6.**
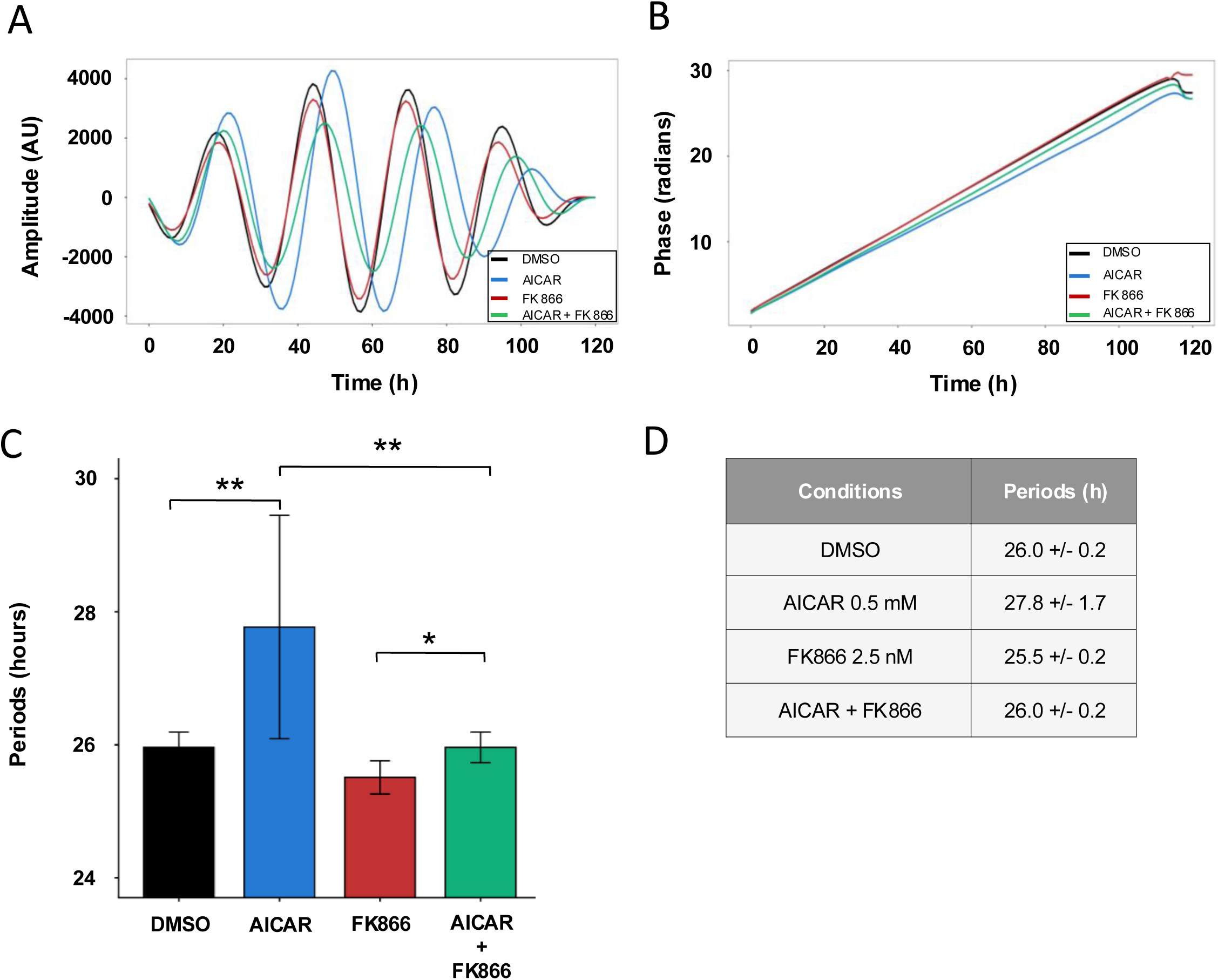
Effect of AICAR with FK866 combination on *Bmal1* promotor activity. A. Filtered signals from U2OS B6 treated with AICAR at 0.5 mM (blue), or FK866 at 2.5 nM (red), or AICAR 0.5 mM + FK866 2.5 nM (green), compared with DMSO (black). B. Instantaneous phases from A. C. Periods were inferred from the instantaneous phases in B during the interval 40-110h of the experiment. Statistical analysis was performed using a Kruskal–Wallis test to assess differences among conditions (***, *p* < 0.001), followed by Wilcoxon-Mann-Whitney test for multiple comparisons, and p-values adjusted using the Holm correction, * p<0.05, **p<0.01. N=3 for 3 independent experiment, n=9. D. Table of periods in hours +/-SD.

### AMPK-driven clock modulation depends on the timing of AICAR delivery

To further test this hypothesis, we then sought to determine how the timing of AICAR administration affects the subsequent phase of the oscillations. For that purpose, we added 0.5mM AICAR to different wells every 4 hours to span a whole circadian period; other wells receiving the same quantity of DMSO as vehicle needed to administer 0.5 mM AICAR. Wells were initially synchronized as in previous experiments. To correlate the administration time with the progression through the circadian cycle, the first AICAR administration was synchronized with the first minimum of the luminescence signal, corresponding in principle with the smallest *Bmal1* transcription rate, as discussed previously.

The conventional way to characterize how the clock reacts to an input at different points of its cycle is to compute a phase response curve (PRC) (Forger 2017), where a transient disturbance is applied during a specific time interval, before returning to the initial condition. The PRC displays the resulting phase shift as a function of administration time, which quantifies the sensitivity of the clock to this input during the given time interval. The PRC thus provides a specific signature of the clock together with its input pathway.

We wondered whether we could establish a PRC for AICAR administration, although determining a PRC is experimentally challenging. The classical method could not be applied because, once AICAR was delivered to cells, they remained continuously exposed to the compound to avoid altering clock phase with a medium change. We therefore compared the asymptotic phases of two wells having received AICAR at different times. Because the two cultures would eventually be exposed to identical conditions, they would converge to the same oscillation frequency, making their phase difference constant over time and thus well defined. Moreover, the histories of the two wells were identical except in the time interval between the two AICAR administrations. The final phase difference between the signals therefore provides a measure of the effect of AMPK activation on p*Bmal1* during this time interval, offering insight into the underlying mechanism. An acceptable time resolution is obtained by comparing wells in which AICAR was administered four hours apart, at times t_n_ = t_0_+ 4n hours to study the action of AMPK in the four-hour intervals [t_n,_t_n+1_] (Fig. 7A). To precisely quantify these results, the phases were determined using the same Hilbert transform method as before (see Material and Methods). For clarity, the phase shifts were converted from radians to ordinary time by considering that 2π radians correspond to one period of 27.5 hours.

**Figure 7.**
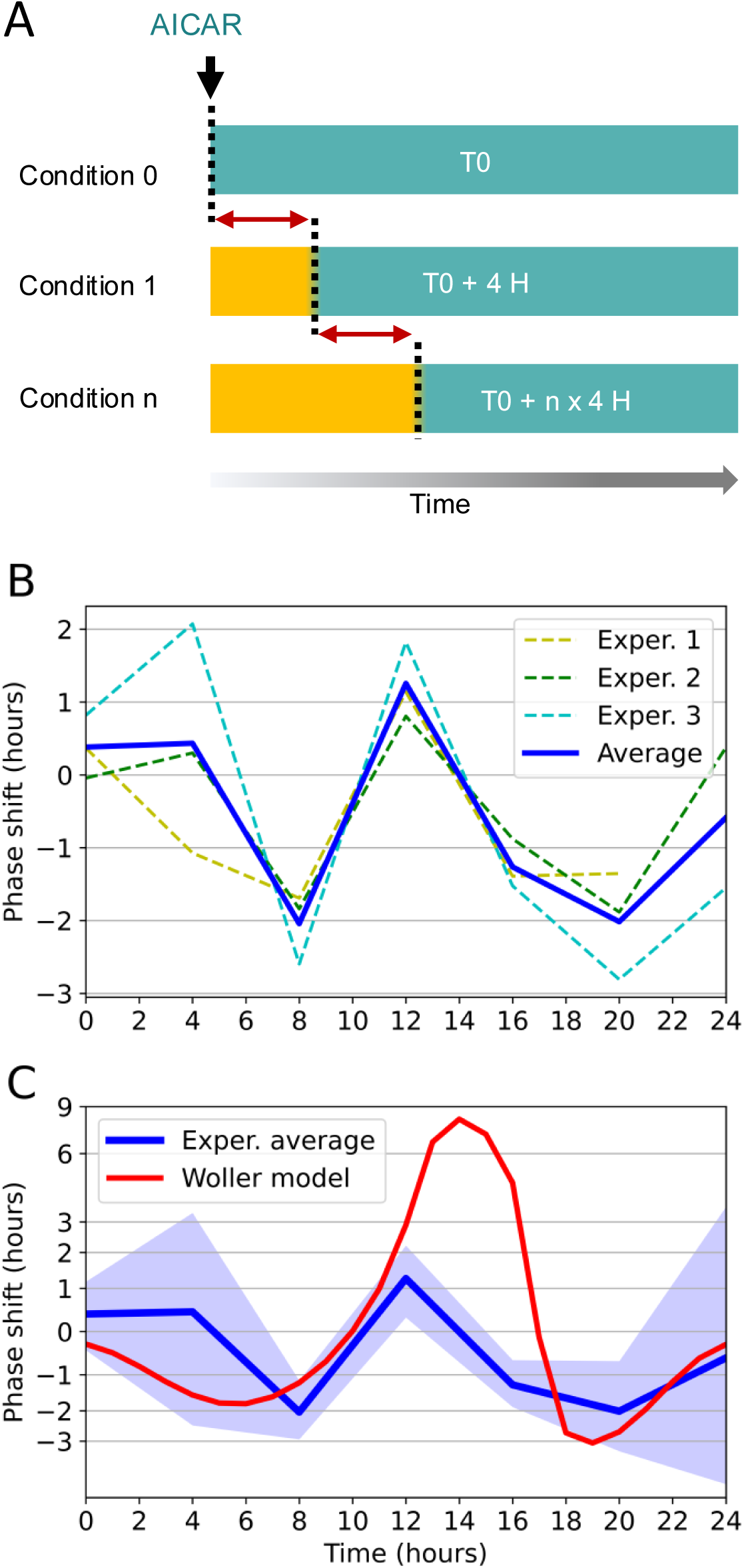
Clock phase response curves to AICAR timely administration. A. Protocol to measure a PRC-like curve without changing medium. The time interval when U2OS cells are exposed to AICAR is represented in green while the yellow color indicates the time interval preceding AICAR administration (marked by a dashed line). By measuring the difference in phase between conditions N and N+1, we can estimate the phase shift induced by exposition to AICAR in the 4-hour time interval indicated by the red double arrow. This phase shift is then corrected by substracting the phase shift observed with DMSO. B The experimental PRC was obtained in three independent experiments. C The theoretical PRC computed from the modified Woller model is compared to the average experimental PRC and its 95% confidence interval, as described in the text, with phase shift expressed in hours. Note the nonlinear scale used for a better display of the experimental PRC.

Interestingly, AICAR administration modified the circadian oscillations depending on time of delivery (Fig. S8). We validated that these effects were not due to DMSO vehicle since its administration at different timepoints did not alter the circadian oscillations (Fig.S8). The resulting phase response curve is shown in Fig. 7B for the three experiments that we conducted to assess the reliability of the measurements, as well as their average.

Although there is variability for some time points, the general structure of the PRC curve is very consistent across the three experiments, with an unusual W (or “mexican hat”) shape where two negative extrema are separated by one positive extremum. Remarkably, the positive extremum is observed at 12 hours, consistent with our previous observation that the maximal effect of AICAR on luminescence level was obtained 12 hours after the beginning of the circadian cycle (see Fig. 2C).

Then, we computed the PRC of the mathematical model used to compute Figs. 3B and 3C. Remarkably, we observed that the theoretical PRC displayed a very similar W shape, and that it fitted reasonably well within a 95% confidence interval around the experimental PRC, except for the 16-hour data point. Between 12 and 16 hours, the theoretical PRC displays a sharp peak associated with a strong phase advance, before returning in the vicinity of the experimental curves at 18 hours (Fig. 7C). In the mathematical model used, we found that this time window corresponds to a strong variation in NAMPT activity, and accordingly in NAD+ abundance and PGC-1-alpha activity. In liver cells, for which the model was designed, *Nampt* is indeed under circadian control whereas it is constitutively expressed in U2OS cells (Hughes et al. 2009, data accessible at NCBI GEO database, accession number GSE13949), which could explain the discrepancy observed.

Given this important physiological difference, the overall consistency between the experimental and theoretical PRCs is quite remarkable and suggests that the model nevertheless captures essential ingredients of the molecular mechanisms involved.

## DISCUSSION

While there is now a relative consensus on the principal actors of the core clock circuitry, the precise mechanisms allowing the hepatic and other peripheral clocks to be entrained to diurnal metabolic cycles are still under investigation, even if some input pathways have been proposed in the literature, such as AMPK-dependent CRY1 destabilization (Lamia et al. 2009) or enhancing of PER translation by insulin (Crosby et al. 2019).

Guided by the theoretical study of Woller et al. (2016), we investigated experimentally and theoretically the response of the U2-OS clock to AMPK activation. A key actor of the Woller mathematical model is the PGC-1-alpha protein, which integrates signals from both AMPK and SIRT1 (Cantó et al. 2010) to regulate *Bmal1* transcription (Fig. 8). The importance of PGC-1-alpha was also noted by Foteinou et al. (2018) who, however, focused exclusively on SIRT1 and did not consider AMPK. In contrast, Woller et al. (2016) proposed that AMPK was the main input. Here we further investigated this hypothesis by combining biological and numerical experiments.

**Figure 8.**
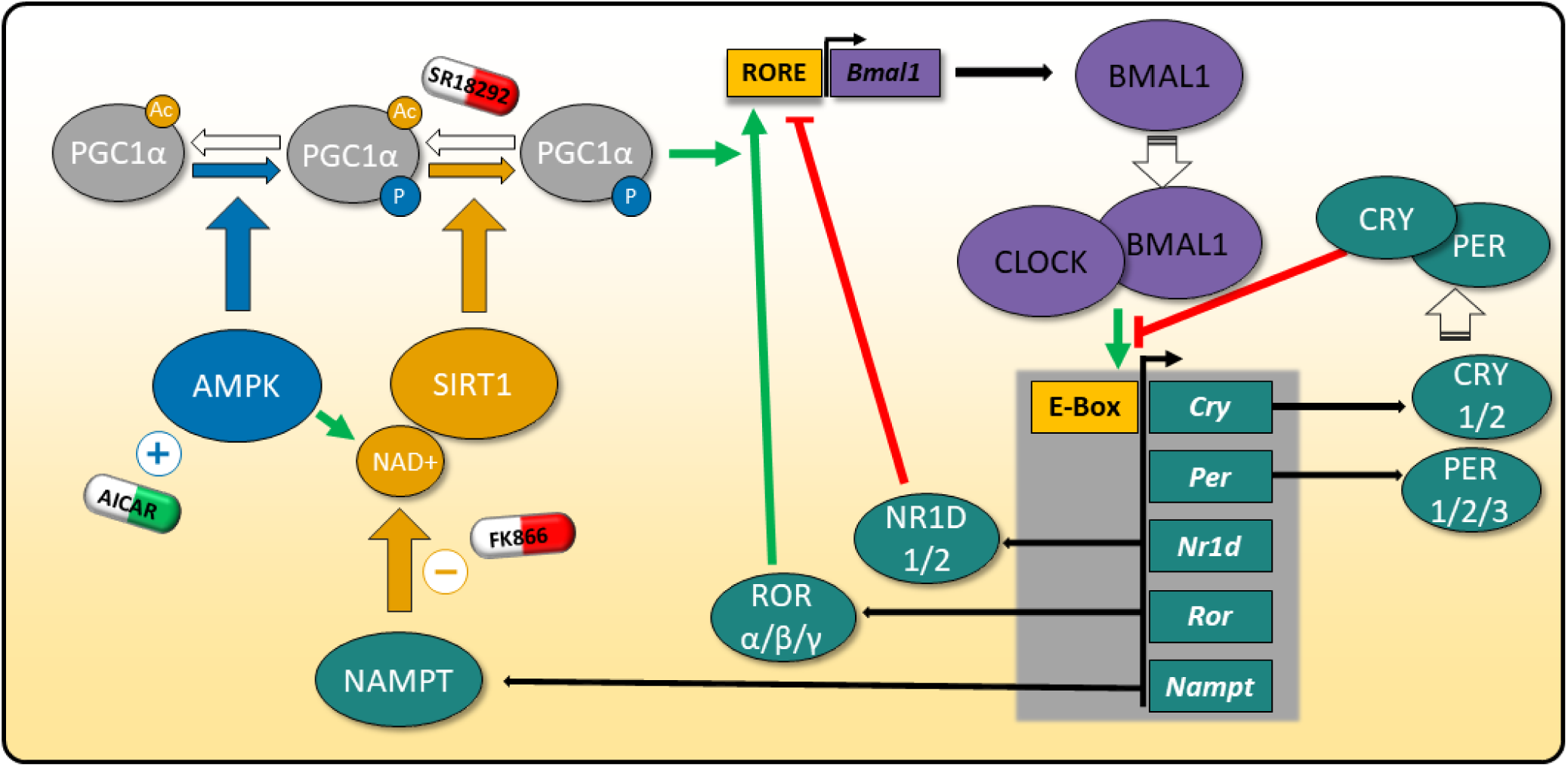
Proposed model recapitulating how AICAR, SR18292 and FK866 impact the circadian clock. Metabolic factors AMPK and SIRT1 influence the feedback loop based on the NAMPT/SIRT1/PGC-1-alpha/BMAL1 axis. AICAR activates AMPK, whereas FK866 inhibits NAMPT, thereby blocking the NAD+ salvage pathway, inducing NAD+ depletion and reducing SIRT1 activity. PGC-1-alpha integrates AMPK and SIRT1 signaling to amplify ROR-dependent activation of *Bmal1* transcription, the AMPK-mediated phosphorylation of PGC-1-alpha being a prerequisite to its deacetylation by SIRT1. SR18292 promotes the interaction of PGC-1-alpha with the acetylase GCN5, reducing PGC-1-alpha activity. Consistent with this mechanism, AICAR increases *Bmal1* promoter activity and lengthens the clock period, while FK866 counteracts these effects. Administration of SR18292 alone shortens the clock period and reduces the amplitude of clock oscillations. When combined with AICAR, however, the effect of SR18292 appears to be overridden, consistent with the ability of AMPK activation to elevate NAD+ levels and thereby sustain SIRT1 activity, counteracting the action of GCN5.

First, we observed that AMPK activation dramatically increased *Bmal1* promoter activity, with an AICAR-driven dose-dependent increase in the luminescence signal of up to 100% when cells are exposed to 0.5 mM AICAR, which was the highest dose preserving long-term viability. Using a biosensor, we verified that this dose, close to AICAR IC_50_ for AMPK (Corton et al. 1995), induces a near-optimal, although not total, AMPK activation. This is consistent with the findings of Lamia et al. (2009), who reported that in mouse liver, several clock gene expression profiles are dampened upon knocking out LKB1, thereby impairing AMPK phosphorylation. In U2OS-B6 cells, however, Lamia et al. (2009) observed that AICAR administration at 2 mM induced a decrease in the average *Bmal1-luc* level. At the same dose, we observed that the U2OS-B6 cell count was strongly diminished over time (Figs. S1A and S1B), and the luminescence signal level was accordingly reduced. This is consistent with reports that AICAR induces mitochondrial apoptosis in human osteosarcoma cells at 1 mM and above (Morishita et al. 2016).

Beyond the effect on expression, we found a gradual lengthening of the circadian period length as the AICAR dose is increased, changing from 25.7 hours in vehicle-treated cells to 27.5 hours for 0.5 mM AICAR. This agrees with the observation of Lamia et al. (2009) that, upon administering 2 mM AICAR, the period increased from 25.3 hours to about 31.3 hours.

We then checked whether our experimental observations could be predicted by the mathematical model of Woller et al. (2016). While this model was originally fitted to experimental data from mouse livers, its mathematical structure only depends on the molecular interaction network. This network is identical in all organs, although the kinetic constants may be tissue dependent. Moreover, the expression profiles of some clock genes, such as *Bmal1*, are remarkably similar in different tissues (see, e.g., (Zhang et al. 2014)), suggesting that a mathematical model designed for one tissue may be useful in the context of another.

A comprehensive sensitivity analysis of the Woller model surprisingly revealed that over the 96 kinetic constants on which the model relies, only the *Bmal1* basal transcription rate and the PGC-1-alpha nuclear abundance had a strong influence on *Bmal1* expression levels. These two parameters directly control *Bmal1* transcription since PGC-1-alpha is a coactivator of *Bmal1*. In particular, modifying the kinetic constants associated with the repressor NR1D1 had almost no effect on *Bmal1* expression, suggesting that the latter is not very sensitive to the duration of the derepression phase, and depends essentially on regulations that are active during this phase. Overall, this sensitivity analysis confirmed that in the Woller model, the co-activation of ROR-family transcription factors by PGC-1-alpha, leading to enhanced *Bmal1* transcription, is the dominant pathway through which AMPK activity influences the clock, despite the model incorporating several other interactions, including AMPK-dependent CRY1 degradation.Interestingly, the sensitivity analysis pointed to a deficiency of the Woller model, which did not take into account the documented self-amplification of nuclear PGC-1-alpha activity, due to its nuclear translocation upon activation (see, e.g., (Anderson et al. 2008; Fan et al. 2020)) or to its autoregulation (Handschin et al. 2003). To take into account this critical molecular control, the model was thus modified to make PGC-1-alpha nuclear abundance AICAR-dependent. Specifically, the basal values and sensitivities to AICAR of AMPK activity and PGC-1-alpha nuclear abundance were four freely adjustable constants while the other parts of the model were left unchanged except for a global time rescaling to match the control period. Numerical simulations of this marginally modified model could simultaneously reproduce the strong increase in transcription and the variation of the period with AICAR dose.

Such agreement is far from trivial given the complexity of the clock model and the extensive interactions among fitted parameters that determine the collective dynamics of the clock. This suggests that although the Woller model was originally developed for hepatocytes, one of its key ingredients - the regulation of *Bmal1* transcription by PGC1-alpha – may operate similarly in the two cell types and could underlie the strong impact of AICAR on *Bmal1* promoter activity observed in our experiments. Further numerical simulations revealed that enhanced *Bmal1* transcription upon AMPK activation and period lengthening are directly related, since the former effect induces higher NR1D1 peak levels, which lengthens the repression phase and thus increases the time interval between *Bmal1* peaks.

Given the putative role of PGC-1-alpha in the mechanism, we tested the effect of the pharmacological PGC-1-alpha inhibitor SR18292 in U2OS-B6 cells. SR18292 reduces PGC-1-alpha activity by promoting interaction with the acetyl transferase GCN5 (Mutlu et al. 2024). At a dose of 20 µM, the clock period was indeed strongly reduced from 25.7 to 22.5 hours, consistent with the hypothesis that an active PGC-1-alpha lengthens the clock period. We also observed a severe attenuation of the amplitude of circadian oscillations in the luminescence signal, suggesting an important role for PGC-1-alpha in clock operation or synchronization.

However, testing whether SR18292 administration could counteract the effect of AICAR did not yield clear evidence, as the inhibitory effect of SR18292 appeared to progressively diminish over time. The fact that AICAR enhances PGC1-alpha deacetylation by stimulating NAD+ abundance and SIRT1 activity (Cantó et al. 2009), thus directly opposing the effect of SR18292, is a plausible explanation of this experimental artefact. This suggested to directly target NAD+.

Thus, we used AICAR in combination with FK866, an inhibitor of NAMPT, a key enzyme of the NAD+ salvage pathway. After the first circadian cycle, the *Bmal1* promoter activity increase and period lengthening induced by AICAR were almost completely abolished by FK866, indicating that the action of AMPK on the circadian clock is strongly NAD+-dependent. The small residual effect may be due to other interactions of AMPK with the circadian clock, including AMPK-dependent CRY1 degradation (Lamia et al. 2009) or PER2 destabilization via CKIepsilon (Um et al. 2007).

This dependence on NAD+ provides an important clue for distinguishing between the possible mechanisms through which AMPK influences the clock. It supports the hypothesis that AMPK acts primarily through PGC-1-alpha, whose activation requires both phosphorylation by AMPK and deacetylation by the NAD+-dependent decetylase SIRT1 (Cantó et al. 2010). It is also consistent with the observation by Foteinou et al. (2018) that silencing *SIRT1* expression reduced *Bmal1:*luc expression by approximately one half.

The PGC-1-alpha hypothesis also explains why the effect of AICAR, measured through the positive variation in the time derivative of the luminescence signals, is undetectable when *Bmal1* promoter activity is at its minimum (Fig. 2), suggesting that AMPK influences the clock through a molecular actor required for *Bmal1* transcription. Indeed, AMPK-activated PGC-1-alpha, which is a potent co-activator of the RORs (Liu et al. 2007), cannot influence *Bmal1* transcription until ROR-family proteins occupy the ROR/REVERB response element (RORE) of the *Bmal1* promoter, otherwise bound by its competitor NR1D1. In a recent analysis of a simple hepatic clock model, Delpierre and Lefranc (2026) likewise identified enhanced *Bmal1* transcription as a potential mechanism through which the variation in a metabolic factor between day and night could entrain the clock.

Changes in period length naturally induce phase shifts and support the role of molecular actors as synchronizing inputs. A more complete characterization of how a given pathway synchronizes the clock can be achieved by measuring a phase response curve (PRC), which provides a signature of both the core clock and its input pathways (Taylor et al. 2008; Forger 2017). To avoid spurious phase resetting caused by medium washing, we used a protocol where AMPK was added irreversibly to the medium at different times in separate cell cultures, starting at the time of minimal promoter activity, and then at 4-hour intervals. We then compared the phase difference between cell cultures treated four hours apart, allowing us to quantify the effect of AICAR in the time interval between administrations. This approach allowed us to characterize the variation in AICAR sensitivity across the circadian cycle, and compare the experimental measurements with mathematical model predictions.

Remarkably, we found that the response curve had an unusual “W” (or “mexican hat”) shape, with two negative minima and one positive maximum, located in the middle of the circadian cycle. Of note, the temporal profiles of NAD+ and AMP measured in mouse livers (Hatori et al. 2012) have two maxima through the diurnal cycle. For comparison, the phase response curve obtained by Crosby et al. (2019) with insulin in liver was a monotonously decreasing curve. However, the AMPK and the insulin pathways may act relatively independently, as suggested by recent experiments showing that knocking out the insulin receptor in mouse liver has little influence on *Bmal1* and *Nr1d1* expression profiles in mice fed *ad libitum* (Fougeray et al. 2022).

In numerical simulations of our mathematical model, the theoretical PRC displayed the same W-shaped profile as the experimental one and fell largely within its 95% confidence interval. The most notable discrepancy occurred at the 16 h timepoint, where the theoretical predictions diverged strongly from the experimental measurement. Few previous works have determined and directly compared experimental and theoretical phase response curves (Taylor et al. 2014; Thommen et al. 2015). The overall agreement observed here is remarkable given that the mathematical model was minimally modified from the one developed for liver, suggesting that it captures important features of the clock dynamics. In fact, the discrepancy observed during the second half of the circadian cycle illustrates the sensitivity of the PRC to details of the underlying mechanisms, as it very likely detects the important physiological differences between hepatocytes and U2OS cells. Importantly, our theoretical results suggest that larger phase shifts may be observed in hepatocyte cells compared to U2OS cells, reinforcing the potential of AMPK as an important modifier of the liver circadian clock, both in amplitude and period. This motivates future experiments to measure the PRC in hepatocytes.

The strong influence of AMPK activation on the expression of the master clock activator *Bmal1* is an important result, as *Bmal1* contributes to the regulation of most other clock genes. Changes in *Bmal1* expression may thus affect the temporal orchestration of many downstream genes. Notably, dampened clock oscillations have been associated with several pathological conditions. In particular, reduced clock gene expression has been observed in mice fed a high-fat diet (Hatori et al. 2012; Eckel-Mahan et al. 2013), with clock disruption preceding metabolic disorders (Eckel-Mahan et al. 2013). Attenuation of circadian oscillations is also a hallmark of aging (Kondratova et Kondratov 2012; Mattis and Sehgal 2016; Hood and Amir 2017) and has been reported in several neurodegenerative disorders (Leng et al. 2019; Fan et al. 2022; Zheng et al. 2023). Moreover, our finding that AMPK action on the clock is strongly NAD+-dependent may provide a link between the decline in NAD+ levels (Covarrubias et al. 2021) and reduced *Bmal1* expression observed in aging.

To conclude, we combined biological and mathematical approaches to provide new evidence about the integration of AMP and NAD+ levels as a key mediator in the pathway leading from metabolic sensing to *Bmal1*, the master clock activator, as commented by Furlan et al. (2019). Our experimental and numerical results were all consistent with the hypothesis that PGC-1-alpha plays a key role in this pathway, as proposed by Woller et al. (2016) and Foteinou et al. (2018). Future experimental studies will aim to dissect the complexity of this mechanism in hepatic cells, especially as diminished amplitudes for several clock genes were reported by Lamia et al. (2009) upon *LKB1* knock-out, consistent with our observations in U2OS-B6 cells. Measuring the PRC in this context could be useful to refine and improve the Woller model.

More generally, a better understanding of the mechanisms governing *Bmal1* expression could have broad consequences for health and disease. In this context, AMPK and PGC-1-alpha could emerge as promising chronotherapeutic targets for manipulating circadian clock amplitude in pathological conditions in which it appears to be a causative factor.

## ACKNOWLEDGEMENTS

We thank all members of our team, GDR CNRS 2588 Imabio and GDR CNRS 2108 AQV for their help and for their fruitful discussion/comments. We are much grateful to Dr DiTacchio for kindly providing us with U20S B6 cells.

## AUTHOR CONTRIBUTIONS

AV, AF and ML designed research. AV, AT, AB, RTI and PL performed experiments. PD and ML carried out numerical simulations. All authors analyzed and interpreted the data. AV, AF and ML wrote the manuscript. All authors commented, revised and accepted the manuscript.

## DATA AVAILABILITY STATEMENT

Data and material are available on request to

## SUPPLEMENTARY DATA CONFLICT OF INTEREST

The author(s) declared no potential conflicts of interest with respect to the research, authorship, and/or publication of this article

## MATERIAL AND METHODS

### Cell culture

U2OS B6 cells bearing the *Bmal1:luc* reporter are a kind gift from Dr Di Tacchio (Vollmers et al. 2008). They express destabilized firefly luciferase (pGL4.22, Promega, Madison, Wisconsin) under the control of *Bmal1* promoter (-422 to +108). They were cultured in Dulbecco’s Modified Eagle’s Medium (DMEM) with a glucose concentration of 4.5 g/L (41966 - Thermofisher), supplemented with sodium pyruvate 1mM, GlutaMAX 1X, 10% fetal bovine serum (FBS), and antibiotics (100 U/ml of penicillin and 100 µg/ml of streptomycin), and selected with puromycin at 2 µg/mL. Cells are maintained at 37°C and 5% CO2 in incubators with a humidified atmosphere. Cell culture is split twice a week to prevent reaching high confluence (>80%). Cells are detached with a 0.25% trypsin solution supplemented with 0.5 mM EDTA, then centrifuged at 200 g for 2 minutes, washed with fresh medium, and seeded at 8,000 cells/cm² in a new flask.

### Pharmacological tools

Pharmacological modulators such as AICAR (Stemcell) or FK866 (Sigma Aldrich) were added to the acquisition media at the start of the experiment to achieve concentrations of 0.1, 0.2, 0.5, 1, and 2 mM of AICAR, and concentrations of 1, 5, 10, or 20 nM of FK866. These drugs were dissolved in DMSO, so cells treated with equivalent volumes of DMSO were used as controls in the experiments. The same amount of the DMSO vehicle was used for AICAR doses from 0 to 0.5 mM, to ensure that differences between curves would only arise from the difference in AICAR dose. 2DG (D317-Merck-Millipore) is added at 10mM final concentration to saturate AMPK activity.

### Viability assessment

The crystal violet assay is performed by starting with a cell rinsing step using PBS. Then, crystal violet is added to the wells containing the cells for 20 minutes at room temperature. The cells are washed with water. Finally, methanol is added to the wells to solubilize the adherent cells at the bottom of the wells after one hour of incubation at room temperature. The absorbance at 570nm is then measured in each well with a plate reader.

### Bioluminescence acquisition

Cells were seeded into 96-well plates at a density of 20,000 cells/well. The day after, they were synchronized by changing the medium from the culture medium to the acquisition medium with a rinsing step using Calcium-and Magnesium-free PBS (Lonza). The acquisition medium was supplemented with 0.2 mM luciferin (Promega E1602, Beetle Luciferin, Potassium Salt), and administered with a final volume of 200 µL per well. Wells at the edges of the plate were not seeded with cells to avoid the edge effect, but filled with sterile water to maintain an optimal humidity in the plate. The plate reader used is a FLUOstar Omega (BMG Labtech) running on Omega 5.11 software.

### Signal processing for estimating the action of AICAR on *Bmal1 transcription*

Luminescence traces were approximated using cubic splines obtained using the splrep function from the interpolate package of the python Scipy v1.10 numerical library. The approximation was excellent given that the traces are very smooth. The second derivatives of the splines were obtained using the splev function from the library, and then integrated to get an estimate of the time derivative of the luminescence signal. This procedure allowed us to elegantly remove linear trends as well as vertical offsets in the luminescence signals due to variations in cell count.

### Signal processing for estimating period

Raw bioluminescence signals were analyzed to assess their periodicity, which served as a criterion for selecting data exhibiting circadian rhythmicity. For this purpose, the “JTK” (Hughes et al. 2010) algorithm implemented in the “Metacycle” library was used. Time series that did not pass the rhythmicity test (Bonferroni FDR>0.05) were excluded of the study.

Then, a signal processing protocol was developed, based on signal filtering in the frequency domain. The signals were handled by a Python script (version 3.8.13) run in the RStudio environment (version 2022.7.1.554) with R version 4.2.1 (R Core Team, 2021). Data were filtered using a second-order Butterworth bandpass filter (Python library: SciPy, functions: butter and lfilter), configured to retain signals with frequencies ranging from 7.71 to 17.36 µHz, corresponding to periods between 16 and 36 hours.

Once the signals were filtered, a Hilbert transform was applied (Python library: SciPy, function: hilbert) to calculate the instantaneous phase ϕ vs time as being the argument of the analytical signal. The time derivative of this phase divided by 2π is the instantaneous frequency *f* of the signal. Thus we estimated the period by carrying out a linear regression on the curve representing the phase, excluding an initial transient, which yielded an average slope α, from which the period T can be obtained according to the following formulas.

The availability of a continuous estimate of the instantaneous frequency is useful as it allows one to check the constancy of this estimate over time and thus the absence of change in the experimental conditions.

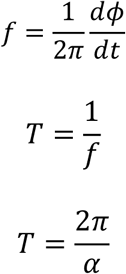

We checked that the DMSO vehicle used to resuspend AICAR did not modify the period when used in different volumes and proportions similar to those used in the AICAR experiments (Fig. S3).

### Phase response curve calculation

PRCs were computed using two different methods

### Hilbert transform method

The instantaneous phases for each curve corresponding to administration at Tn are obtained with the Hilbert transform as described previously. The phase difference between the T_n_ and T_n+1_ conditions is computed with the following formula where the effect of DMSO is subtracted from the effect of AICAR: Δφ=(φAICAR(Tn)-φDMSO(Tn))-(φAICAR(Tn+1)-φDMSO(Tn+1)), where φAICAR(Tn) (resp., φDMSO(Tn)) denotes the asymptotic phase of the culture treated with AICAR (resp., DMSO alone) at time Tn, measured at a given common time. When this quantity is positive, it implies that the well treated at time Tn displays a phase advance as a result of the exposition of AICAR between Tn and Tn+1 Several PRC curves are calculated and averaged from T=80h to T=110h (experiment time) that correspond to the last part of experiment.

### Difference of phase base on circadian peaks methods

The peaks of every circadian cycle are reported and the difference of time between equivalent peaks (circadian cycle number 2,3 or 4). First, the difference of time between AICAR and DMSO condition is made then the PRC is calculated by subtracting Tn and Tn+1 condition. The strategy is the same as described above, but the “phase” here corresponds to the experiment time.

### AMPK biosensor measurement

To measure AMPK activity in U2OS B6 cells, we used a plasmid encoding a FRET-based biosensor sensitive on AMPK activity, namely AMPKAR-EV (Konagaya et al., 2017).

U2OS B6 cells were seeded at a density of 100,000 cells in a 35 mm dish with culture medium and transfected at day 1 with 500 ng of the AMPKAR-EV plasmid. At day 2, cells were detached and seeded into µSlide 0.8 chips (ibidi) at a density of 40,000 cells per chip. At day 3, the cells were observed in Fluorobrite medium (Thermo Fisher A1896701) supplemented with 1% sodium pyruvate, 1% glutamax 100X, 100 U/ml penicillin, 100 µg/ml streptomycin, and 10% FBS. The acquisition started 1 hour after culture stabilization in the microscope incubation chamber set at 37°C. AMPK modulators such as AICAR and 2-DG were prepared at appropriate concentrations and introduced into the reservoirs of the microfluidic station (ARIA - Fluigent). The flow rate was set to 10 µl/min to minimize mechanical stress on the cells, which could affect fluorescence signals.

Fluorescence imaging was performed on a Leica DMI6000B coupled with Quantem 512SC camera (Photometrics). Cells were observed with x20 objective (numerical aperture: 0.5). The illumination source was SOLA U-nIR (lumencore). The whole system was under Micro-Manager control (version 1.4.24) (Edelstein et al. 2010).

### Fluorescence analysis

The background noise of each picture was measured by generating a region of interest in a cell-free zone using Fiji (Schindelin et al. 2012) and then subtracted. Cell segmentation was performed using Cellpose, a deep learning-based cell segmentation tool (Stringer et al. 2021), and cell tracking was then carried out using Trackmate (Tinevez et al. 2017; Ershov et al. 2022). The data generated by Trackmate were processed using RStudio (version 2022.7.1.554) with R. The data presented in graphs are means with standard deviation (SD) calculated for each data point.

### Statistics

First, a Shapiro-Wilk test (shapiro.test) was applied to determine if the datasets follow a normal distribution. Then, a Bartlett test (bartlett.test) was used to compare the variances between conditions. If the datasets passed both tests, we performed a One-Way ANOVA analysis and Tukey test for multiple comparison. If the dataset didn’t pass Shapiro test, Kruskal-Wallis (kruskal.test) analysis and Wilcoxon-Mann-Whitney (wilcox_test) test for multiple comparison were applied. All statistical tests were made in R with stats and rstatix libraries.

