## Supplemental material (Table+figures) for "Experimental and mathematical models reveal that AMPK modulates circadian clock gene expression and period through NAD+-dependent regulation of *Bmal1*"

**Supplementary Table S1. Dependence on AICAR dose of the two kinetic constants of the Woller model characterizing AMPK activity and PGC-1-alpha nuclear abundance.** The kinetic constant names follow those used in the supplementary material of Woller et al. (2016). Given an AICAR dose $d$ expressed in mM, the value of kinetic constant $K$ is given by : $K = \text{base} + d \times \text{sensitivity}$. These represent the only modifications to the model described by Woller et al. (2016), the dependence of the nuclear PGC-1-alpha on AICAR dose describing the nuclear translocation associated with PGC-1-alpha activation.

| Kinetic constant | Meaning | base | sensitivity |
| --- | --- | --- | --- |
| Act_AMPK | AMPK activity | 0.04505648133 | 7.42681835987 |
| Prot_PGC1a | Nuclear PGC-1-alpha abundance | 1.26559020396 | 2.84102692814 |

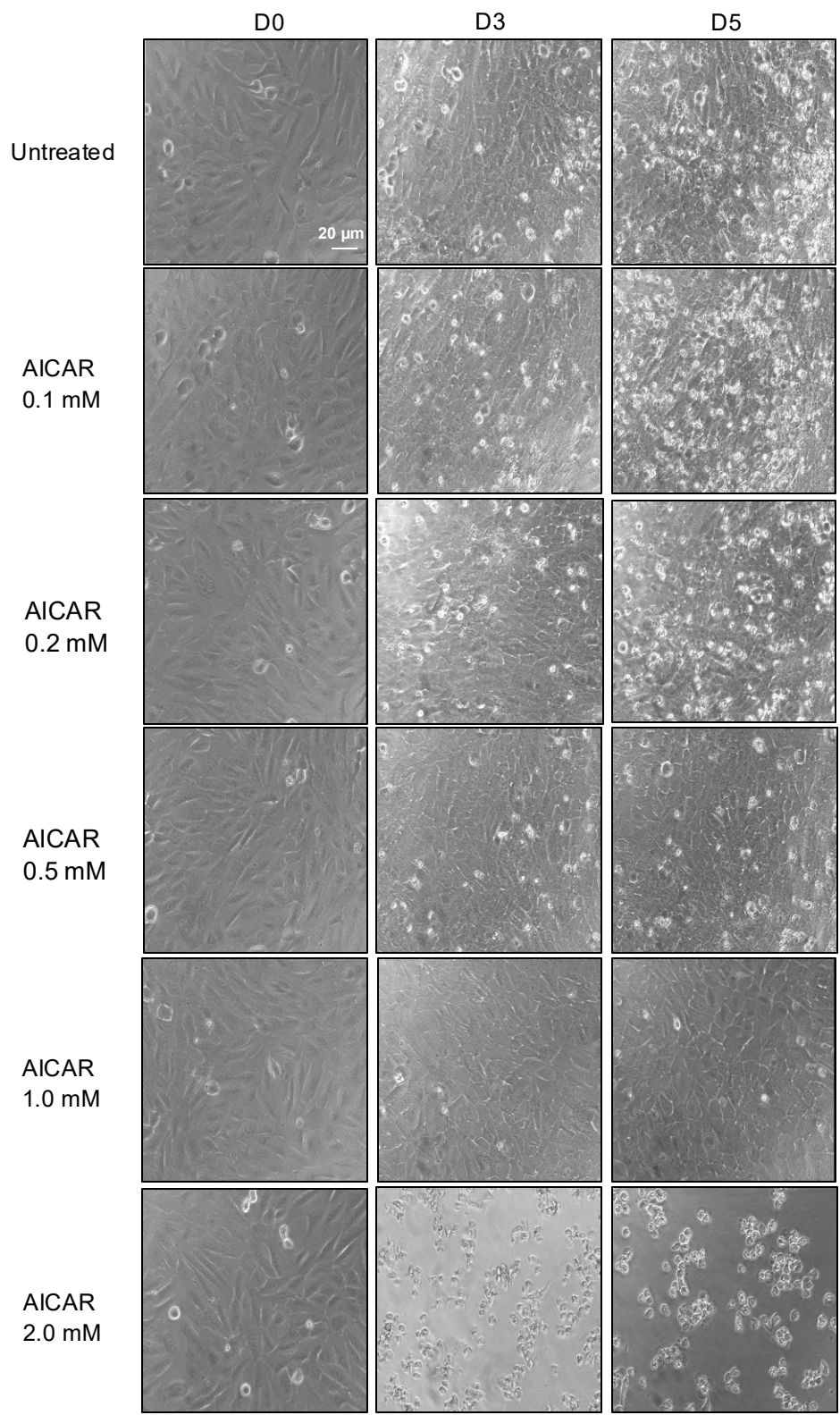

**Figure S1. Phase contrast images of U2OS B6 cells treated with AICAR.** Pictures of cells treated with AICAR at 0.1 mM, 0.2 mM, 0.5 mM, 1 mM and 2 mM over time at day 0 (D0), day 3 (D3) and day 5 (D5). Scale bar 20 μm.

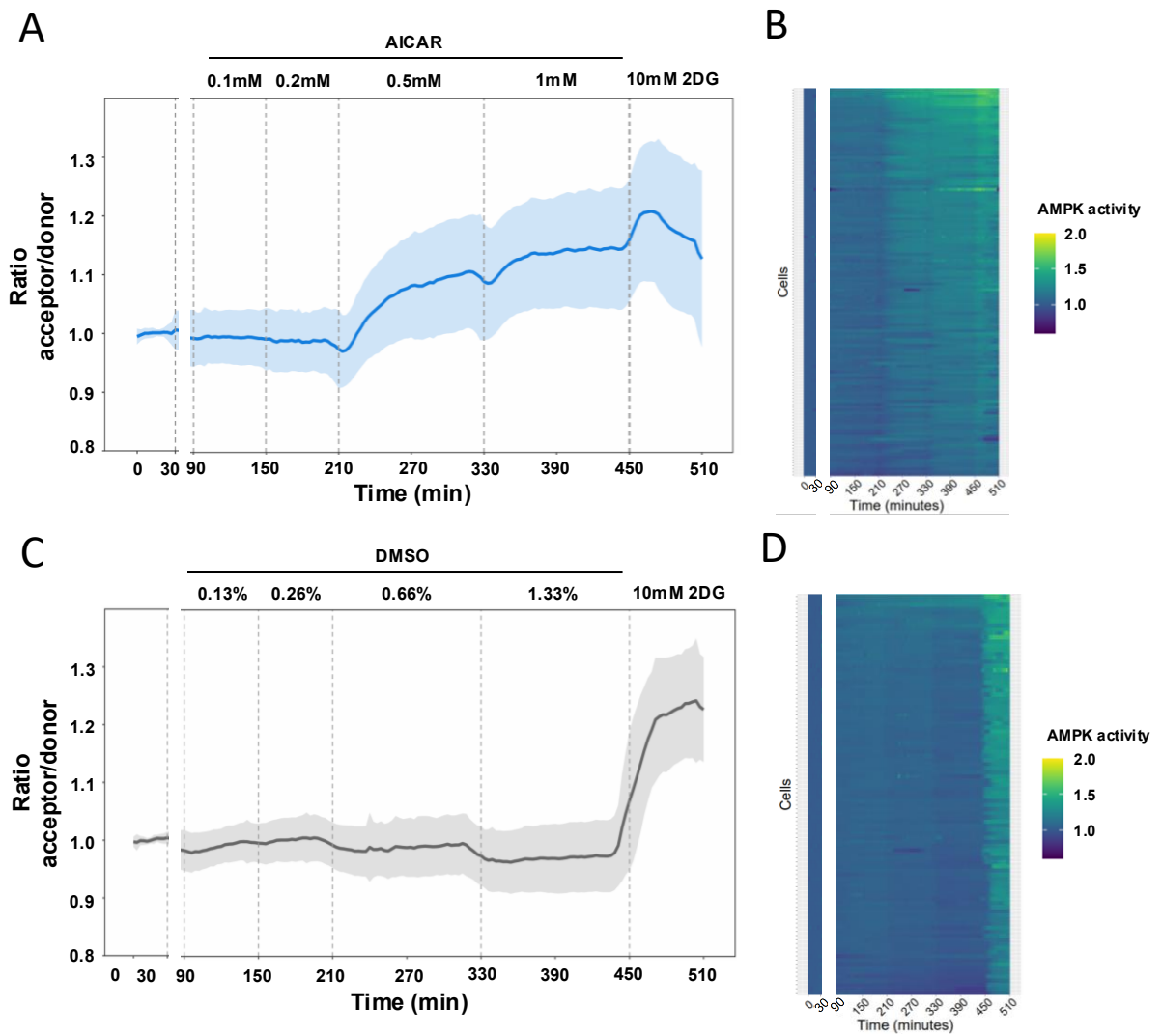

**Figure S2. Measurement of AMPK activity thanks to AMPKAR-EV biosensor.** U2OS B6 cells were transfected with the FRET AMPKAR-EV plasmid (Konagaya et al). AICAR or DMSO used as the vehicle were administered at different concentrations with a microfluidics system. The flux was set to 10 μL/min. **A.** Average (+/- SD) of signal (ratio acceptor/donor) measured in single cells treated with 0.1, 0.2, 0.5 and 1 mM of AICAR, and finally treated with 10 mM 2DG. N=3 independent experiments, n=132 cells. **B.** Kymograph, every line corresponds to single cell treated with AICAR. The flux data are removed (grey dashed bar) and the data after 90min are corrected by flux substractions. **C.** Average (+/- SD) of signals measured in single cells treated with 0.13%, 0.26%, 0.66%, 1.33% DMSO diluted in the medium and finally treated with 10 mM 2DG, N=3 independent experiment, n=98 cells. **D.** Kymograph, every line corresponds to single cell treated with DMSO.

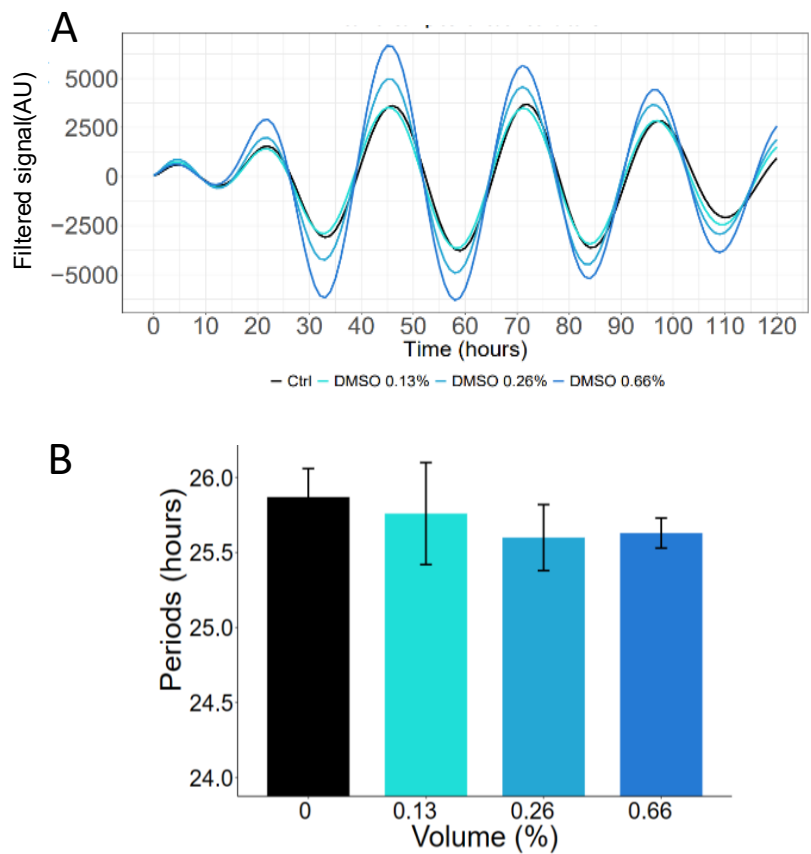

**Figure S3. Measurement of clock periods in response to DMSO administration.** U2OS B6 cells were administered with increasing concentrations of DMSO used as the vehicle (for AICAR treatments in parallel). A. Filtered signals are provided. B. Periods were inferred from treated signals.

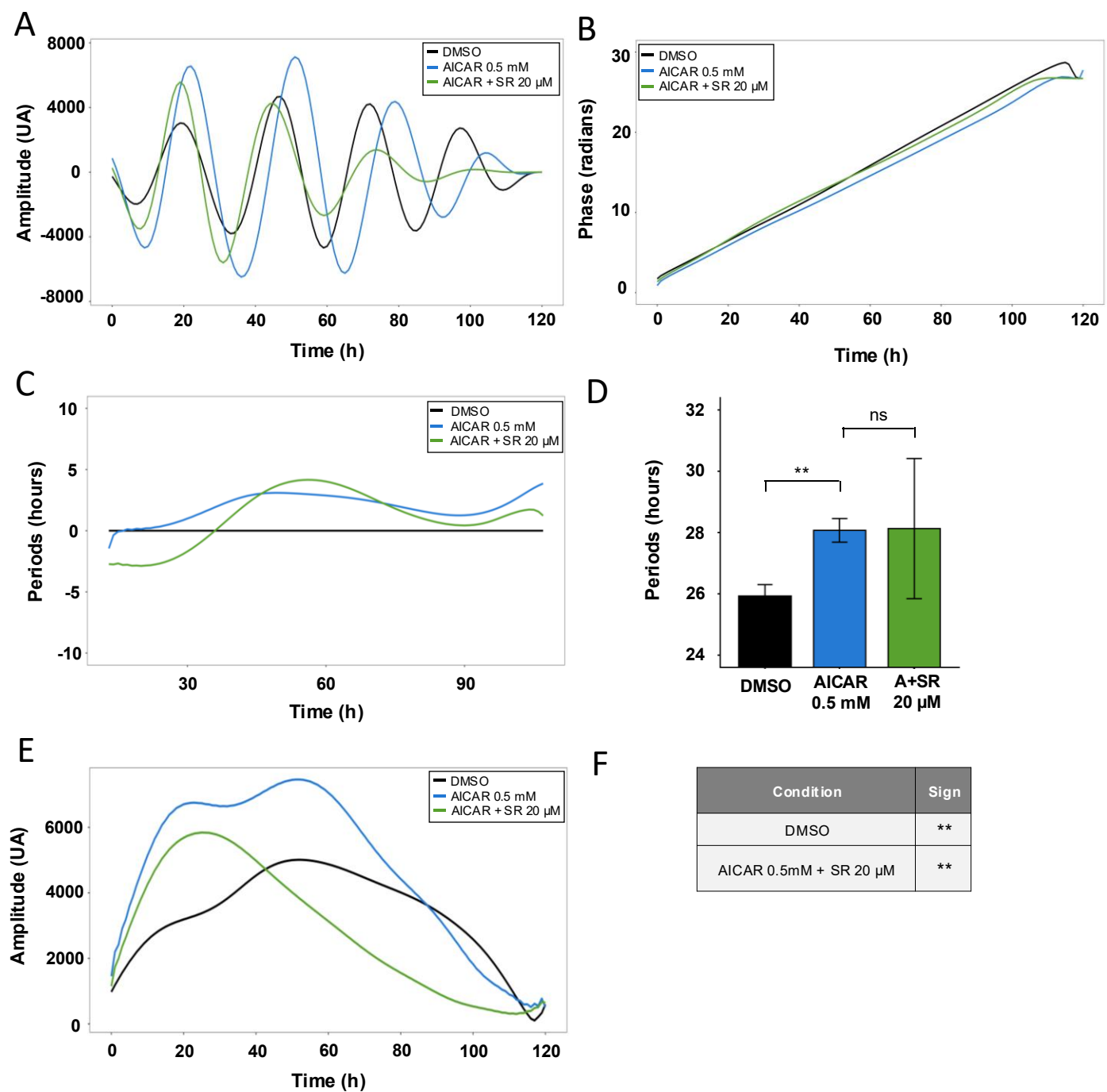

**Figure S4. Effect of the combination of AICAR with SR-18292 on *Bmal1* promoter activity.** A. Filtered signals processed using a Butterworth filter are shown in blue for AICAR alone, in green for treatment with SR-18292 at 20 μM combined with AICAR at 0.5 mM, and in black for DMSO control. B. Instantaneous phases calculated from A using Hilbert transform. C. Instantaneous periods, calculated from B and averaged over 26h time windows centered on axis time, relative to DMSO condition. D. Periods, in hours ± SD, were inferred from the instantaneous phases in B during the interval 40-100h of the experiment for DMSO (25.6 ± 0.2); AICAR 0.5 mM (28.1 ± 0.4) and AICAR 0.5 mM + SR 20 μM (28.1 ± 2.3). Statistical tests were performed with Wilcoxon-Mann-Whitney test for comparisons, and p-values adjusted using the Holm correction, ns : non significant, \*\* : $p < 0.01$. N=3 for 3 independent experiments, n=9. E. Instantaneous magnitude calculated from A using Hilbert transform. F. Comparison of area under curves (AUC) calculated from E, performed with Wilcoxon-Mann-Whitney test for comparisons with AICAR 0.5 mM ; p-values are adjusted using the Holm correction, ns : non significant, \*\* : $p < 0.01$. N=3 for 3 independent experiments, n=9.

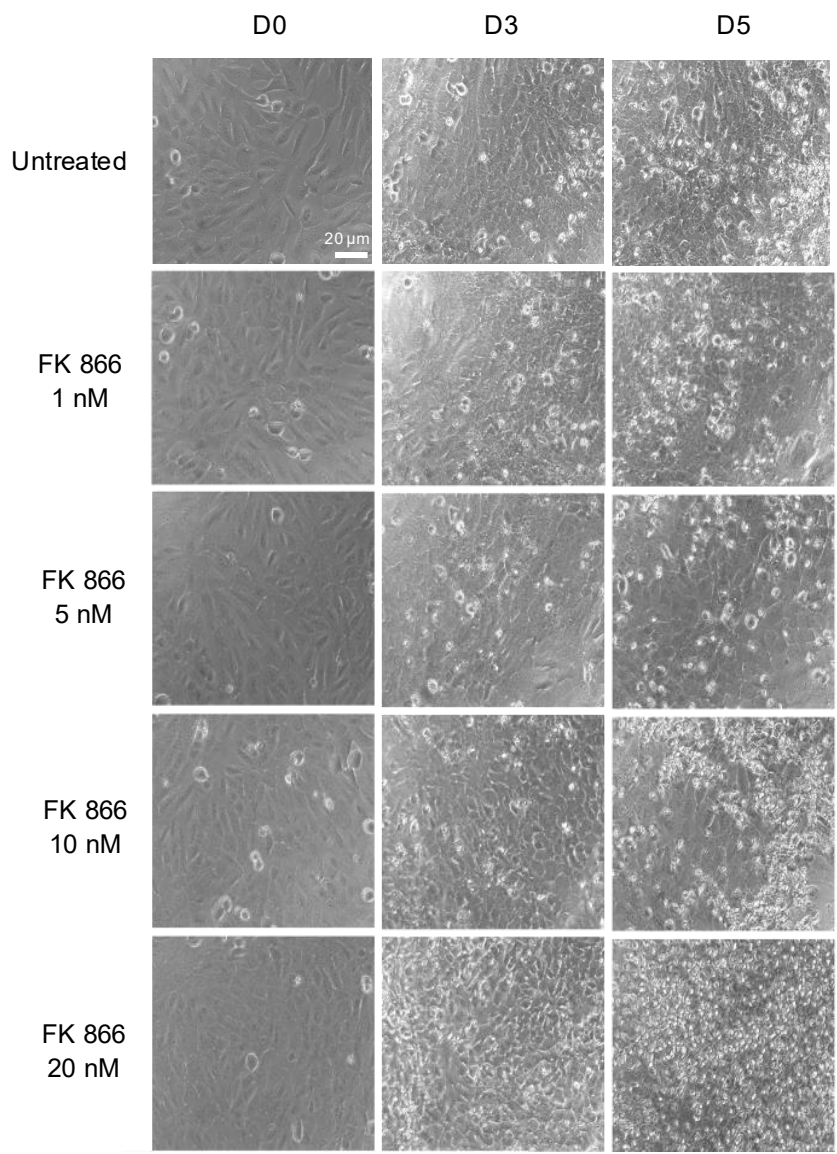

**Figure S5. Phase contrast images of U2OS B6 cells treated with FK866.** Pictures of cells treated with FK 866 at 1 nM, 5 nM, 10 nM and 20 nM over time at day 0 (D0), day 3 (D3) and day 5 (D5). Scale bar 20 μm.

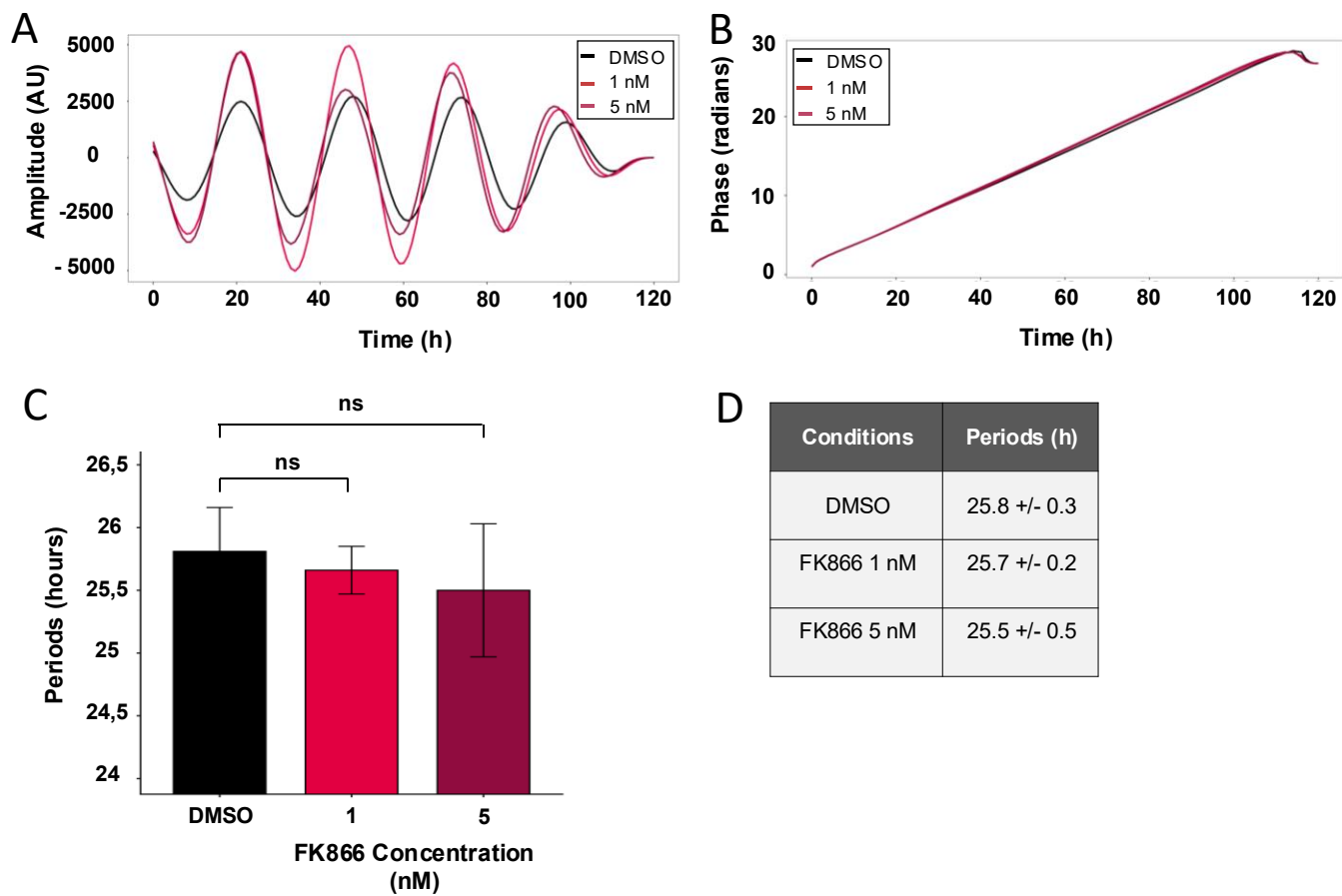

**Figure S6. Effect of FK866 on *Bmal1* promoter activity.** A. Signals were treated with Butterworth filter. DMSO control cells are represented in black and FK866-treated (1nM and 5nM) cells in red. B. Instantaneous phases calculated by using Hilbert transformed from A. C. Periods were inferred from the instantaneous phases in B during interval 40-110h of the experiment. Statistical tests were performed with Wilcoxon-Mann-Whitney test for comparisons, and p-values adjusted using the Holm correction, ns: non significant. D. Table of periods in hours +/- SD.

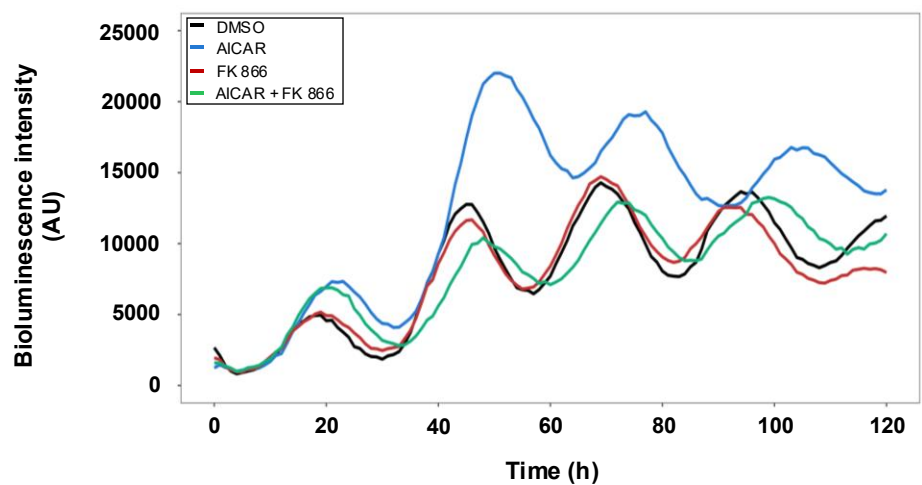

**Figure S7. Effect of combination of AICAR with FK866 on U2OS B6 cells.** Mean of raw signals of U2OS B6 treated with AICAR at 0.5mM (blue), FK 866 at 2.5 nM (red), or combination of AICAR 0.5 mM with FK866 2.5nM (green).

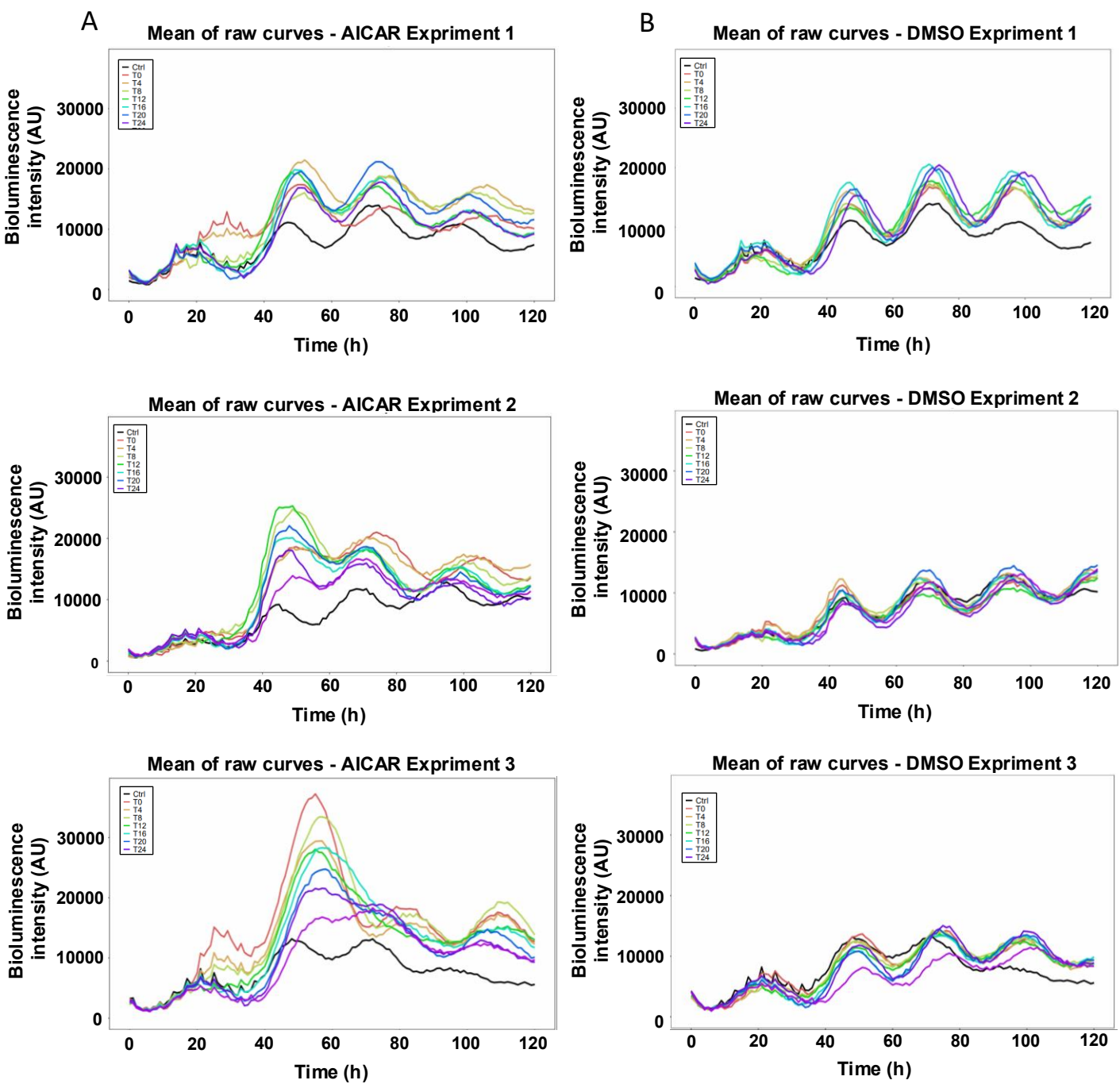

**Figure S8. Raw signals for each PRC experiments:** Mean of raw luminescent signals from U2OS B6 cells treated every 4 hours, from T0 to T0+28h, with either AICAR 0.5 mM or an equivalent quantity of DMSO. N=3 independent experiments.
